# Design of full-length agonist and antagonist antibodies for hard targets

**DOI:** 10.64898/2026.10.21.760573

**Authors:** Robin A. Aglietti, Ramin Ansari, Peter M. Bowers, Tim-Henrik Buelles, Hadrien Chauvin, Thibault M. Duplay, Mohamed El Hibouri, Julien Jorda, Joshua Laniado, Matthias M. A. Malago, Kyle Meador, Anya Polovina, Katty Polyak, Justyna Sosna, Robert Tacke, Todd O. Yeates, Lucas Zanini

## Abstract

Therapeutic antibodies have transformed modern medicine, yet conventional discovery relies on lengthy experimental workflows, offers limited control over how antibodies engage their targets, and struggles to deliver candidates across many important target classes. Despite recent advances in computational protein design, generating full-length antibodies with desired functional activity and favorable developability profiles remains challenging. Here we present an end-to-end computational design pipeline built around Pythia-1, an all-atom generative model that reasons jointly over amino acid sequence and three-dimensional atomic structure under explicit design constraints. We generated candidates in full-length human immunoglobulin G (IgG) format across 22 therapeutic targets spanning receptor ectodomains, cytokines, growth factors, and G protein-coupled receptors (GPCRs), a particularly challenging target class for antibody discovery. Following a single computational design round, experimental screening of at most 48 candidates per target identified antibodies with single-digit nanomolar affinities for soluble proteins and single-digit nanomolar cell-binding EC_50_ values for GPCRs. The pipeline’s ability to target specified epitopes and conformational states enabled the design of antibodies with agonist and antagonist activity. *In vitro* characterization demonstrated target-specific engagement and potent functional modulation, with single-digit nanomolar inhibitory potency in functional assays for a range of targets, including multiple GPCRs. To our knowledge, this study provides the first demonstration of both GPCR agonism and antagonism by *de novo* designed full-length IgG antibodies. We incorporated key developability and manufacturability requirements into the initial designs by conditioning generation on human germline frameworks and applying sequence constraints to limit known liabilities. The resulting antibody designs exhibited biophysical profiles comparable to or better than those of clinical-stage therapeutic controls. We provide the complete sequences of all lead candidates to support independent experimental validation and further development. Together, these findings demonstrate that our technology can deliver antibodies with programmable function and key properties expected of lead candidates, opening opportunities for first-in-class therapeutics against hard-to-drug targets.

## 1 Introduction

Monoclonal antibodies (mAbs) have become one of the most successful therapeutic modalities, with applications across many areas of medicine [1, 2]. Antibodies offer several advantages over small-molecule drugs, including target specificity, favorable pharmacokinetic properties, and the capacity for multivalent and multispecific binding. More than 200 antibody drugs have been approved globally [3], with over 1,400 undergoing clinical evaluation [4]. In 2024, global sales of antibody-based therapeutics were estimated at approximately US$267 billion [4], and mAbs accounted for the majority of the ten best-selling drugs worldwide as of 2018 [1]. Despite this success, conventional antibody discovery has struggled to deliver therapeutic candidates across several important target classes, particularly integral membrane proteins such as G protein-coupled receptors (GPCRs), ion channels, and transporters.

Traditional antibody discovery relies on animal immunization and extensive library screening. These approaches offer limited control over whether the identified antibodies combine the functional activity and biophysical properties required for a therapeutic program. Immunization can fail to elicit antibodies against therapeutically relevant epitopes, particularly when those sites are poorly immunogenic, conformationally dynamic, or difficult to access on membrane proteins with limited extracellular surface area [5–7]. Screening antibody libraries using display technologies selects for target binding, but does not necessarily favor the molecular properties required for therapeutic development. Antibodies identified through phage display, for example, can exhibit polyspecificity, self-interaction, and other unfavorable biophysical characteristics [8]. Extensive optimization is often needed after initial hit discovery. Both immunization and library screening involve lengthy experimental workflows and substantial costs. These limitations have motivated efforts to design antibodies computationally.

Recent advances in deep-learning-based protein design have enabled high experimental success rates for small protein binders, including miniproteins with relatively rigid binding surfaces formed by α-helices and β-strands [9–11]. Designing full-length antibodies with paired heavy and light chains remains more challenging because antigen recognition is mediated primarily by six complementarity-determining region (CDR) loops, three from each chain, whose flexibility and varied backbone geometries make their antigen-bound conformations more difficult to predict and design [12, 13]. Some studies have simplified this problem by designing single-domain antibodies, such as VHHs (nanobodies), which contain three CDRs and no paired light chain [14, 15]. Recent VHH and single domain designs also exploit framework-mediated contacts, which can facilitate interface design by drawing on the framework’s rigid, well-defined secondary structure rather than relying on less structurally predictable CDR loops alone [16]. These framework contacts may, however, introduce additional optimization and developability challenges. As isolated domains, VHHs also lack the Fc region that contributes to the prolonged circulating half-life and effector functions of full-length immunoglobulin G (IgG) antibodies [17].

Several recent studies have reported promising results in the computational design of antibodies against predefined epitopes on soluble proteins and receptor ectodomains [11, 16, 18–20]. Progress toward multipass membrane proteins, particularly GPCRs, has been more limited, with reported functional designs largely using VHH-based formats [11, 21, 22]. Although GPCR binding has also been demonstrated with designs in full-length IgG format [11, 20], to our knowledge, functional modulation of GPCR signaling in cells has not yet been demonstrated for *de novo* computationally designed full-length IgG antibodies. Considerable opportunity therefore remains to design antibodies with defined functional activity and specificity against challenging therapeutic targets.

Here we present an antibody design platform for generating full-length IgGs with defined functional activity and properties suited for therapeutic development. At its core is a protein design pipeline powered by Pythia-1, an all-atom generative model. The platform provides control over where an antibody binds and the conformational state of the antigen it recognizes, allowing the intended mechanism of action to shape the design from the outset. Human framework selection and antibody-specific sequence constraints bring key developability and manufacturability considerations into the initial design to reduce the need for extensive downstream engineering. We benchmark our platform across a broad panel of soluble and membrane-associated targets, including cytokines and receptor ectodomains, and extend the approach to GPCRs by targeting functionally relevant epitopes in active and inactive receptor conformations. Through rapid experimental screening and characterization of full-length IgG designs, we demonstrate high hit rates, high affinity, favorable developability profiles, and functional receptor modulation.

## 2 Results

### 2.1 Generative Pipeline for Protein Design

We have developed a generative pipeline built around Pythia-1, an all-atom generative model trained across multiple tasks on the largest protein structure dataset reported to date. Pythia-1 jointly generates protein backbone geometry, amino acid sequence, and side-chain conformations conditioned on a target structure and specified design constraints. The pipeline additionally incorporates a dedicated sequence-generation model that can optionally generate sequence variants for each structure. Candidates undergo multiple stages of filtering and prioritization, combining model-derived confidence scores, physics-based structural plausibility metrics, and sequence-based metrics (Fig. 1a). The resulting end-to-end computational workflow enables the generation and prioritization of candidates for a target within one day.

**Figure 1.**
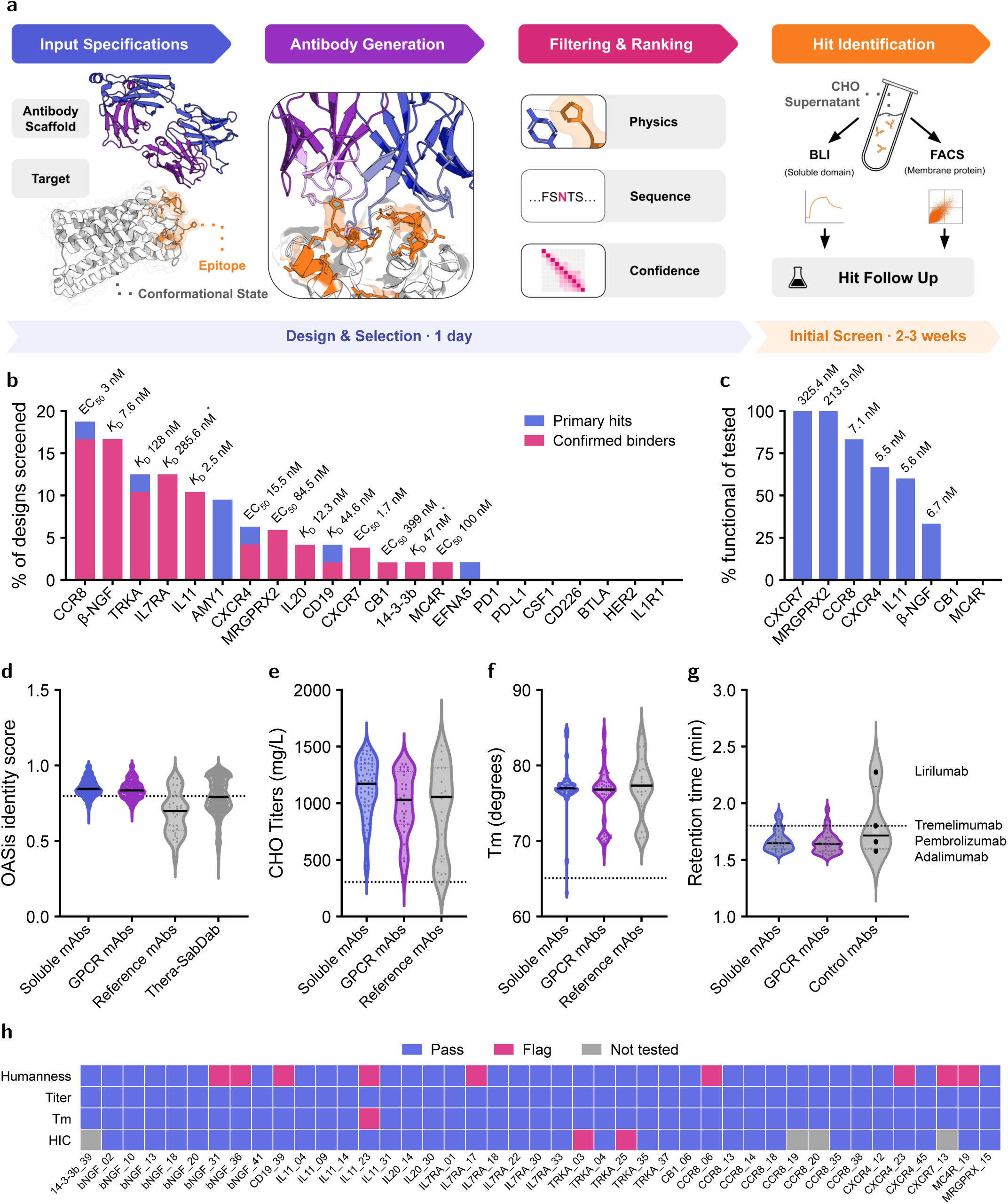
Overview of full-length IgG1 ***de novo*** design on fully human frameworks. (a) Workflow of end-to-end design and screening of full-length IgG1 antibodies, beginning with input specifications of human germline scaffold and structure, target antigen structure, and desired epitope. Conditioned on these inputs, the model generates antibody candidates, which are then filtered and ranked. Final designs are screened as full-length IgGs by applying CHO supernatants to either ProA BLI tips (in the case of soluble target antigens such as cytokines or extracellular domains), or to cells expressing the target protein for flow cytometric analysis (in the case of membrane target proteins). (b) Experimental validation of antibody designs across diverse targets. Antibodies were screened by HTP-BLI for soluble protein and ectodomain targets or by flow cytometry for binding to cell lines overexpressing the indicated GPCR (with the exception of CD226, which was screened by flow cytometry). Primary hits are shown in blue. The criterion for identifying a designed antibody as an initial hit was a total BLI response greater than 0.06 nm at 150 s after antigen association for soluble targets, or a z-score greater than 10 for GPCR targets. Confirmed binders (pink) were defined as antibodies with a *K*_D_ less than 5 µM in a five-point BLI assay for soluble targets, or an EC_50_ less than 1 µM for GPCR targets. The lowest *K*_D_ (soluble targets) or cell-binding EC_50_ (GPCR targets) for each target is indicated above the corresponding bar, with an asterisk denoting values derived from BLI sensorgrams with non-ideal fits to a 1:1 Langmuir model. Targets for which no primary hits were identified are shown with zero primary hit rate. (c) Percentage of tested antibodies exhibiting functional activity for each target. Where activity was observed, the potency IC_50_ (or EC_50_ for MRGPRX2) of the most potent antibody is indicated above the corresponding bar. (d–g) Humanness, expression titer, thermal stability and hydrophobicity were assessed using the OASis identity score, transient CHO expression, DSF and HIC, respectively. In each case, antibodies targeted to soluble targets (blue) and GPCR targets (purple) are compared to reference antibodies against the same set of selected targets. In (d), designs are further compared to all clinically relevant antibodies in Thera-SabDab (grey). In (g), pembrolizumab, adalimumab, tremelimumab, and lirilumab were included as HIC retention-time controls. Dashed lines represent typical standards for each metric based on industry best-practices for clinical candidates, with developability flags below for humanness, titer, and *T*_m_, and above the dashed line for hydrophobic retention by HIC. (h) A heatmap summarizing developability properties of all antibodies with confirmed binding, across humanness (pass *≥* 0.8), transient CHO expression titer (pass *≥* 250 mg/L), Fab melting temperature (*T*_m_) (pass *≥* 65 *^◦^*C), and HIC retention time (pass *≤* tremelimumab at 1.8 min). Each measurement was classified as pass (blue), flag (pink), or not tested (grey) according to the defined developability criteria.

### 2.2 A Platform for Lead-Candidate Antibodies

Building on our pipeline, we set out to develop an antibody design platform specialized for generating candidates with properties suitable for therapeutic development. All designs and subsequent experimental studies reported here used a full-length human IgG1 format. Sequence information from human antibody databases [23] was used to define the allowable design space with respect to CDR-length distributions and to preserve residues at highly conserved positions in framework-adjacent regions. During design, framework regions were constrained to germline sequences from human heavy and light V-gene pairs with known structures and established biophysical properties [24, 25]. CDR sequence constraints were also applied to limit potential sequence liabilities such as N-linked glycosylation and deamidation-prone motifs (Methods, Section 5.3). Designing full-length IgG sequences enabled seamless progression from screening of culture supernatants to affinity measurements, biophysical characterization, and functional characterization of lead candidates.

We selected a total of 15 soluble and ectodomain targets and 7 GPCR targets based on their established therapeutic value, structural diversity, reagent availability and evaluation in prior computational antibody-design studies [11, 14, 26, 27]. For each target, we chose structure coordinate sets and surface epitopes for antibody engagement according to specific functional goals, particularly with respect to whether antagonist and agonist effects were intended. We allocated 360 Pythia-1 designs per soluble and ectodomain target and 720 per GPCR target, with three additional sequence variants generated per design (Methods, Section 5.3), yielding up to 1,440 and 2,880 unique sequences, respectively. After self-consistency and quality filtering, we selected at most 48 designs per target, maximizing structural diversity by prioritizing candidates derived from distinct target-engagement conformations or poses (Supplementary Fig. S8).

For all targets, experimental screening began with production of full-length IgG designs in 1-mL CHO cell cultures. The time interval from DNA synthesis to screening antibody-enriched supernatants was less than three weeks. For soluble/ectodomain targets, antibodies from enriched supernatants were captured on anti-Fc biosensors for BLI, then exposed to media containing soluble, non-avid antigen at a concentration of 1 µM. For GPCR targets, IgG binding to cells expressing the target protein was assessed by flow cytometry using appropriate positive and negative antibody controls and target-negative cells. Primary hits for each target class were produced in 30-mL CHO cell cultures and purified using Protein A beads. Specific, saturable binding was confirmed using five-concentration BLI or dose-dependent cell-surface binding measurements. Promising candidates were characterized in biophysical developability assays, and higher-affinity designs were further evaluated using cell-based functional assays.

### 2.3 IgG Designs Bind Target Antigens With High Affinity

We observed binding of designed IgG antibodies to a broad range of soluble and GPCR therapeutic targets. Cell-surface binding was also evaluated for tropomyosin receptor kinase A (TrkA) and interleukin-7 receptor subunit α (IL7RA) (Supplementary Fig. S10). Primary binding hits, defined by BLI response greater than 0.06 nm or a flow cytometry z-score greater than 10, included 31 hits to 8 of 15 soluble protein targets (53%) and 18 hits across all seven GPCR targets (100%) (Fig. 1b; Supplementary Figs. S1,S3). Affinity characterization showed that 12 of the 31 hits to soluble targets had dissociation constants (*K*_D_) in the single- to double-digit nanomolar range (Supplementary Fig. S2a). Importantly, the designs demonstrated strong specificity, with none of the antibodies showing measurable binding to a negative control antigen (Supplementary Fig. S2b). Likewise, the majority of hits to GPCR targets showed specific binding to transfected cells, with multiple designs exhibiting single-digit nM EC_50_ values (Supplementary Figs. S3,S11). Among the soluble targets, we prioritized interleukin-11 (IL11) and β-nerve growth factor (β-NGF) for functional characterization based on the relatively large number of hits in our primary screen, established functional cell-based assays and the availability of reliable positive controls (Fig. 1c; Supplementary Fig. S2). All GPCR designs with confirmed binding were tested for functional activity, with the exception of C-C chemokine receptor type 8 (CCR8), for which only 6 of the most promising sequences were tested (Supplementary Fig. S3). Individual examples of high-affinity IgG designs illustrating these structure-function relationships are explored below.

### 2.4 Pythia-1 IgG Designs Possess Favorable Developability Properties

We evaluated whether Pythia-1 IgG designs possess biophysical properties appropriate for manufacturing and development. Our analysis included both experimental and bioinformatic analyses of designs, assessing attributes such as production titer, melting temperature in IgG format, humanness, isoelectric point (pI), hydrophobicity, self-association, and other factors that predict formulation and manufacturing success. The majority of designs surpassed acceptable developability thresholds and compared favorably with a panel of clinical-stage and FDA-approved antibodies (Fig. 1d–h; Supplementary Fig. S4a–f) [28, 29]. These results are consistent with the design constraints, including the use of selected V-genes and their corresponding germline framework sequences, and CDR lengths and amino acid compositions representative of native human repertoires [23]. All designs, including those established as confirmed hits, exhibited heavy and light chain CDR amino acid compositions similar to those found in therapeutic antibodies (Supplementary Fig. S5) [28]. The heavy-chain CDR3 length distribution was shifted toward longer sequences relative to that of the Thera-SabDab dataset, particularly among GPCR design hits, which may reflect the requirement to engage recessed binding pockets (Supplementary Fig. S6). Finally, designs demonstrated high structural diversity (Supplementary Fig. S8) and substantial sequence and structure novelty compared to known antibodies cataloged in the Structural Antibody Database (SAbDab) [29] (Supplementary Fig. S7).

### 2.5 Functional IgG Designs for Soluble Therapeutic Targets

We assessed the platform’s ability to generate functional antibodies against a set of soluble protein targets and receptor ectodomains. We selected epitopes where antibody binding was expected to inhibit target function by disrupting interactions with natural binding partners and used Pythia-1 to design antibodies against these sites. This rational approach aims to maximize inhibitory activity while allowing us to vary epitope location and binding orientation to explore structure-function relationships. Our results with antibodies designed against a few selected soluble protein targets are described below.

#### 2.5.1 IL11

Multiple antibodies were obtained against IL11, an interleukin therapeutic target with a potential role in pulmonary and cardiovascular fibrosis [30, 31]. The designed antibodies were targeted to surface regions involved in receptor binding. Initial BLI screening identified 5 hits among 48 IL11 candidate designs with dissociation constants ranging from single- to triple-digit nanomolar values, as determined by follow-up five-concentration BLI analysis (Fig. 2c; Supplementary Fig. S2a). The tightest binder, IL11_04 at 2.5 nM, demonstrated a stronger affinity than the patent-derived IL-11 reference antibody 3C6 (measured at 9.4 nM in the same assay conditions) [32]. To assess cell-based function, we selected 5 of the candidates (*K*_D_ values from 2.5 nM to 504 nM) to test for their ability to disrupt IL11-mediated activation of the IL11R pathway in a reporter assay. The top design, IL11_04, showed potent inhibition of IL11-mediated signaling, with an IC_50_ value of 5.6 nM compared to an IC_50_ of 24.6 nM for the positive control, 3C6 (Fig. 2d). Antibody IL11_23, with *K*_D_=52.8 nM, demonstrated an IC_50_ value of 316 nM. Accordingly, inspection of the design model shows that the surface region on IL11 where the IL11_04 antibody binds overlaps substantially with the natural IL11Rα epitope of IL11 (Fig. 2a,b). The predicted overlap with the IL11Rα-binding surface is consistent with inhibition through receptor competition. IL11_04’s higher affinity may contribute to its greater inhibitory potency, although differences in epitope and binding geometry may also contribute [33]. The weaker affinity of IL11_23 likely contributes to its lower inhibitory potency while the predicted epitope has greater overlap with IL-6Rβ, which binds to the IL11R signaling complex after association of IL11Rα [34].

**Figure 2.**
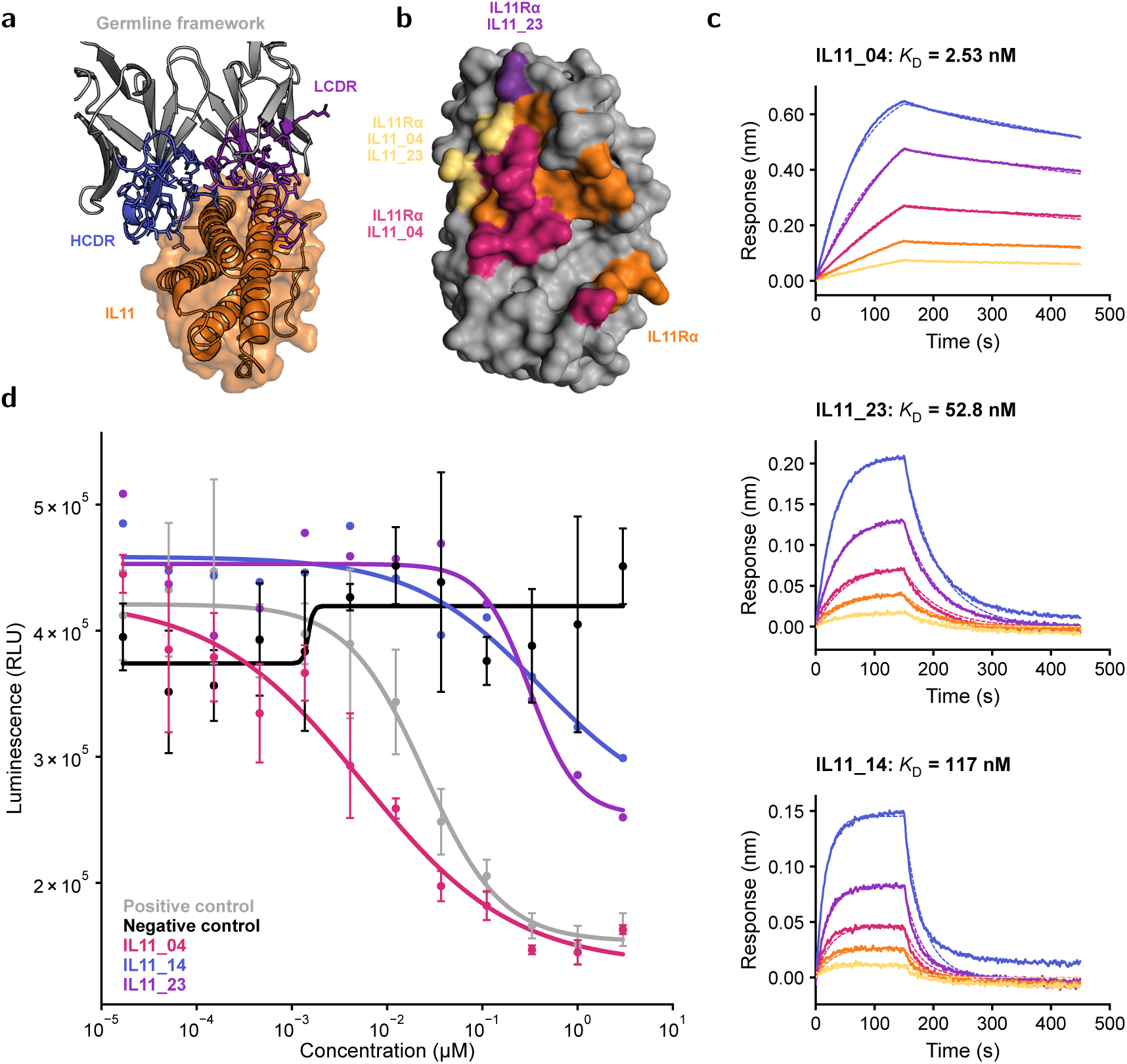
Characterization of designs to IL11. (a) Pythia-1 designed interaction between antibody IL11_04 and IL11. The antibody variable-domain framework is shown in grey, HCDRs and LCDRs in blue and purple respectively, and IL11 in orange. (b) Overlap of selected epitopes with the IL11Rα-binding surface (PDB 8DPS). The IL11Rα epitope includes all non-grey colors, with contacts only to IL11Rα shown in orange. Epitopes overlapping IL11Rα for the anti-IL11_04 model are in pink, while IL11_23 is in purple. Yellow residues are common for all three, IL11Rα, IL11_04, and IL11_23. (c) Binding was assessed by BLI analysis of anti-IL11 antibodies against recombinant IL11. Representative sensorgrams show concentration-dependent binding of IL11 monomeric antigen at five 2-fold serial dilutions to immobilized antibody. IL11_04 and IL11_23 (top, middle) were tested with antigen titrations starting at 50 nM, while IL11_14 (bottom) was tested starting at 100 nM. Data were globally fit using a monomeric 1:1 binding model (dashed lines) to determine the indicated affinities. (d) Anti-IL11 designs were tested for antagonist activity in dose response using the PathHunter® Human IL11 (IL11RA) Signaling STAT3 Reporter Stable Cell Line Assay (HepG2) in the presence of IL11 at a concentration corresponding to EC_80_. The positive control sequence is represented in grey; negative control in black; IL11_04 in pink; IL11_14 in blue; IL11_23 in purple.

#### 2.5.2 β-NGF/TrkA

As a further illustration, we targeted the β-NGF/TrkA signaling pathway, which modulates neuronal survival and differentiation and is relevant to pain management and other therapeutic areas [35–37]. β-NGF forms a two-fold symmetric homodimer that engages two TrkA receptors to promote receptor activation. We targeted the TrkA-binding surfaces of β-NGF with antibodies designed to sterically block engagement with the TrkA receptor and inhibit signaling. Among 48 design candidates tested as unpurified supernatants, 8 anti-β-NGF candidates designated as primary hits were produced, purified, and confirmed to bind β-NGF, with 6 demonstrating single- or double-digit nanomolar affinities using dimeric antigen in five-concentration BLI experiments (Fig. 3b; Supplementary Fig. S2a; Supplementary Fig. S9). Two high-affinity designs, βNGF_18 and βNGF_41, with *K*_D_ values of 7.6 nM and 24.8 nM, respectively, were evaluated for functional activity in a cell-based reporter assay and a TF-1 cell-proliferation assay.

**Figure 3.**
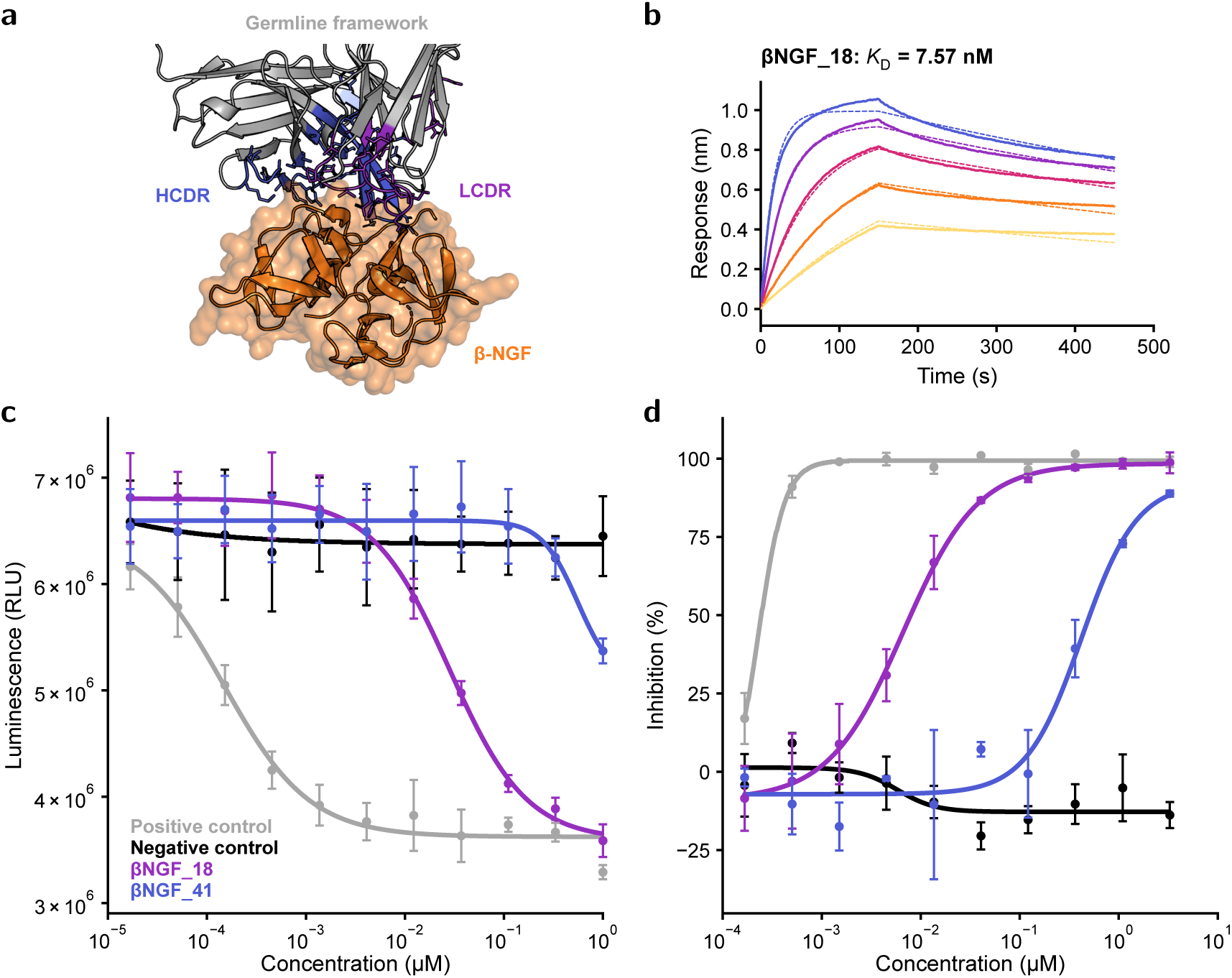
Characterization of designs to β-NGF. (a) Pythia-1-designed interaction between antibody βNGF_18 and β-NGF. The antibody variable-domain framework is shown in grey, HCDRs and LCDRs in blue and purple, respectively, and β-NGF in orange. (b) BLI binding analysis of the anti-β-NGF antibody βNGF_18 against recombinant β-NGF. Representative sensorgrams show concentration-dependent binding of β-NGF to immobilized antibody, using five twofold serial dilutions of antigen starting at 500 nM. Data were globally fit using a 1:1 binding model (dashed lines). Because β-NGF is a homodimer, multivalent interactions may modestly influence the measured binding kinetics and fitted affinity; the reported affinity should therefore be interpreted as an apparent value under these assay conditions. (c) Proliferation of TF-1 cells in the presence of β-NGF and dose-dependent inhibition by anti-β-NGF antibodies were measured using CellTiter-Glo 2.0 after 40 h of incubation. (d) Anti-β-NGF designs were tested for dose-dependent antagonist activity using the luminescent LeadHunter® TRKA Human RTK Kinase Cell Based Antagonist Assay, with β-NGF at a concentration corresponding to EC_80_. In (c) and (d), black denotes the negative control, grey the positive control, purple βNGF_18, and blue βNGF_41.

Both assays report on β-NGF-mediated TrkA activation through downstream signaling and cell activation. The highest-affinity antibody, βNGF_18, exhibited potent antagonism with an IC_50_ value of 6.7 nM in the cell reporter assay (Fig. 3d) and 28 nM in the TF-1 cell-proliferation assay (Fig. 3c), but had significantly lower inhibitory potency than the higher-affinity control antibody [38, 39]. The other selected design, βNGF_41, showed weaker activity in both assays. 6 primary hits were obtained against TrkA at its β-NGF-binding site. 5, with modest affinities ranging from 128 nM to 3.1 µM (Supplementary Fig. S2a; Supplementary Fig. S9), were evaluated in the TF-1 cell-proliferation assay but did not show inhibitory activity.

### 2.6 Functional IgG Designs for GPCR Targets

We assessed the binding and function of IgG designs directed against diverse GPCRs, targeting epitopes from structures exhibiting antagonist or agonist-biased conformations. Across the set of GPCRs, several antibodies were obtained showing clear functional activity in cell-based assays. An analysis of functionally active designs highlights features relevant to the performance of our generative pipeline. We identify designs that exhibit single-digit nanomolar inhibitory potency in functional assays across multiple GPCR targets. We find cases in which the light chain, absent from single-domain antibodies, is predicted to play a critical role in GPCR binding. We also observe, based on predicted models, cases in which antibody CDR loops extend deep into GPCR binding cavities. In a particularly notable case, we demonstrate agonist activity for the target receptor, as discussed below.

#### 2.6.1 Antibodies Designed to Antagonize CCR8

CCR8 is highly expressed on activated, tumor-infiltrating regulatory T cells (Tregs), where CCL1-CCR8 signaling supports recruitment, survival, and immunosuppressive activity [40–42]. Antibodies were designed to antagonize CCR8 based on the structure of its inactive conformation (PDB 8TLM [43]). While this PDB structure was present in Pythia-1’s training set, the maximum CDR sequence identity to the resulting designs was 41% with a minimum CDR edit distance of 40 (data not shown). Binding of antibody design candidates to CCR8 was assessed by flow cytometry using CCR8-expressing cells, with primary hits further evaluated using purified antibodies in flow cytometry cell binding assays. Design candidate CCR8_38 exhibited an EC_50_ of 3 nM, and three other antibodies had measured EC_50_ values below 10 nM (Fig. 4b; Supplementary Fig. S11). All candidates demonstrated target specificity based on binding magnitude and apparent EC_50_ relative to off-target controls, with most showing no detectable binding to HEK cells expressing the unrelated GPCR Mas-related G protein-coupled receptor member X2 (MRGPRX2) (Fig. 4b). The predicted models of CCR8_38 and several other structurally similar variants were notable for the extensive interactions their LCDR1 loops make within the orthosteric pocket of CCR8, facilitated by LCDR1 loops that were 18 residues in length, slightly longer than typically observed [23].

**Figure 4.**
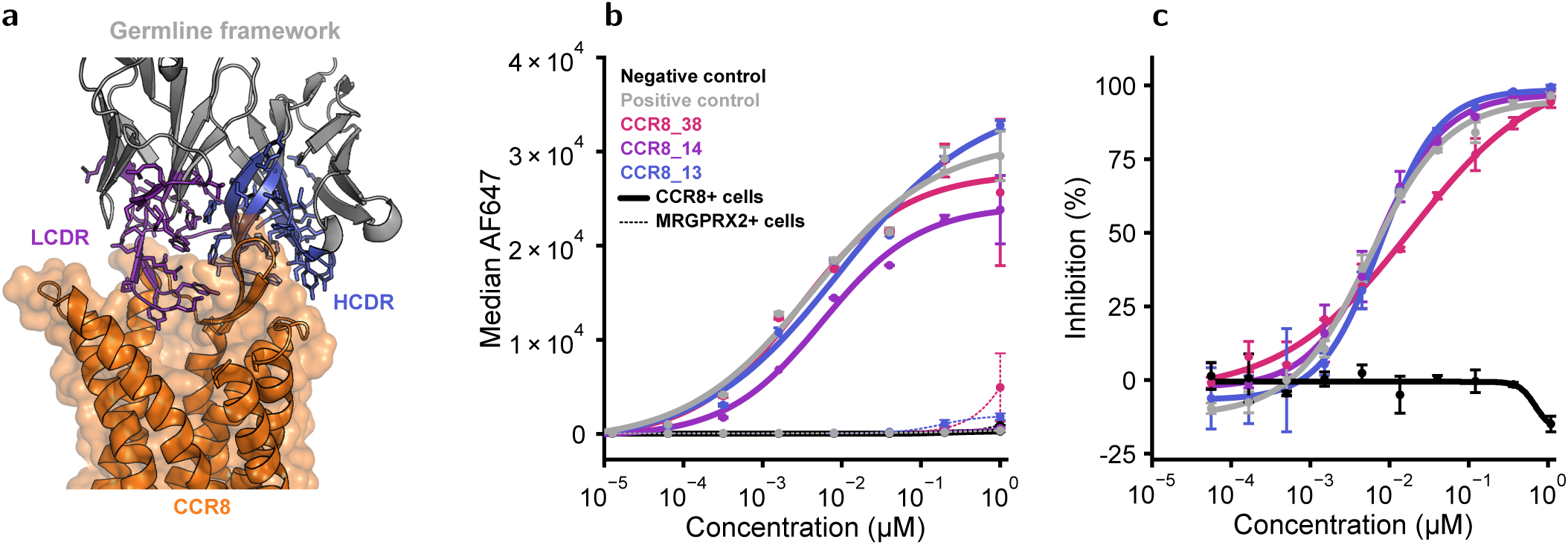
Characterization of designs to CCR8. (a) The predicted CCR8_38 complex shows contacts involving HC and LC CDR1 loops. (b) Concentration-dependent binding of selected antibody sequences to CCR8-overexpressing cells measured by flow cytometry. Binding responses were fit using nonlinear regression to determine on-cell EC_50_ values (solid lines). Binding of the same antibody sequences to MRGPRX2-overexpressing cells (dashed lines) was evaluated as an off-target cellular specificity control. (c) Functional antagonism of CCR8 signaling was assessed using the CCR8 PathHunter® β-Arrestin recruitment assay. CCR8 signaling was stimulated with CCL1 at a concentration corresponding to EC_80_. Antibody concentration-response curves are shown as percent inhibition of CCL1-induced β-arrestin recruitment. Curves were fit by nonlinear regression to determine functional inhibitory potency. In panels (b) and (c), black curves represent negative control; grey curves are positive controls; blue curve is CCR8_13; purple curve is CCR8_14; pink curve is CCR8_38.

We tested this set of high-affinity candidates for their functional effects on CCR8-expressing cells. Antibodies were evaluated for their ability to antagonize CCR8 by blocking CCL1-induced β-arrestin recruitment (Fig. 4c). 5 antibodies showed potent antagonism; 2 (CCR8_13 and CCR8_14) had IC_50_ values in the single-digit nanomolar range. Interestingly, across these candidates, binding EC_50_ values did not correlate strongly with reporter-assay IC_50_ values; the ratios between paired values (IC_50_/EC_50_) ranged from approximately 0.8 to 6.3 (Supplementary Fig. S11). Differences observed here between binding and functional activity emphasize the importance of assessing IgG-formatted candidates in functional studies.

#### 2.6.2 Antibodies Designed to Antagonize CXCR4

Screening identified 2 antibodies that bound C-X-C chemokine receptor type 4 (CXCR4), a GPCR that mediates immune-cell trafficking, hematopoietic stem-cell retention, and tissue repair, making it an important target for stem-cell mobilization, B cell development and inflammatory disorders [44–46]. CXCR4_23 had the lowest apparent cell-binding EC_50_ among the designed CXCR4 antibodies, with an EC_50_ of 15.5 nM (Fig. 5b). We complemented cell binding studies with cell-based functional assays. In a β-arrestin recruitment assay, CXCR4_23 showed an IC_50_ of 5.5 nM (Fig. 5c). The published antibody REGN7663 [47], used as a positive comparator, had an EC_50_ of 1.1 nM but weaker inhibitory activity, with an IC_50_ of 33 nM. Despite its higher apparent cell-binding EC_50_, CXCR4_23 had an approximately sixfold lower functional IC_50_ than REGN7663 in this assay.

**Figure 5.**
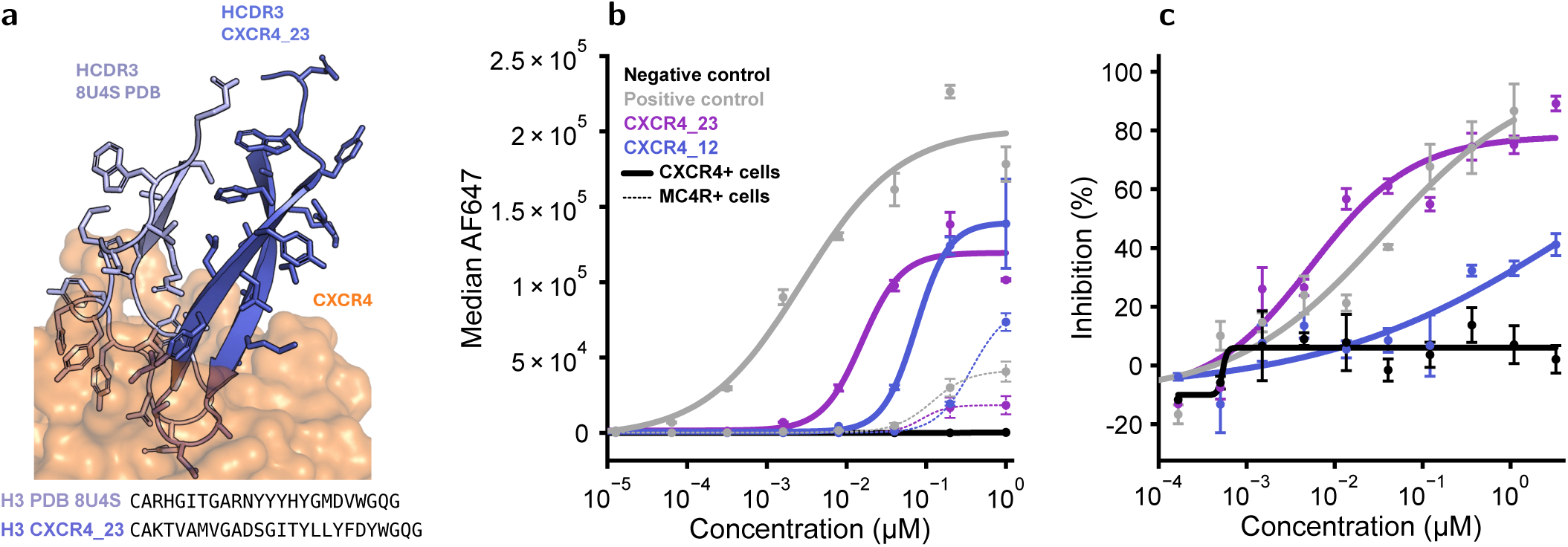
Characterization of designs to CXCR4. (a) Comparison of the HCDR3 loop from the model of designed antibody CXCR4_23 vs a previously published antibody complex structure (PDB 8U4S [47]), highlighting the distinct poses and penetration depth of each antibody. Structures are aligned to CXCR4 in each complex. CXCR4_23 HCDR3 is colored in dark blue, PDB 8U4S anti-CXCR4 HCDR3 in light blue, and CXCR4 in orange. The corresponding HCDR3 sequences are shown below. (b) Concentration-dependent binding of selected antibody sequences to CXCR4-overexpressing cells measured by flow cytometry. Binding responses were fit using nonlinear regression to determine on-cell EC_50_ values (solid lines). Binding of the same antibody sequences to MC4R-overexpressing cells was evaluated as an off-target cellular specificity control (dashed lines). (c) Functional antagonism of CXCR4 signaling was assessed using the CXCR4 PathHunter® β-Arrestin recruitment assay. CXCR4 signaling was stimulated with an EC_80_ concentration of CXCL12, and antibody concentration-response curves are shown as percent inhibition of CXCL12-induced β-arrestin recruitment. Curves were fit by nonlinear regression to determine functional inhibitory potency. In panels (b) and (c), negative control represented as black; positive control as grey; CXCR4_12 as blue; CXCR4_23 as purple.

CXCR4 antibodies obtained in this campaign were notable for the depth their HCDR3 loops reach in the recessed orthosteric pocket (Fig. 5a). The CXCR4 cavity binds its native ligand, CXCL12, with its N terminus extended deeply into the central cavity. The apparent advantage of accessing GPCR binding cavities with smaller nanobodies, unencumbered by a second variable domain, has been noted [48, 49]. Indeed, while multiple such cases have been reported for nanobodies [50–52], CDR penetration into the orthosteric pocket by IgG antibodies is rare. The published IgG antibody REGN7663, obtained by traditional experimental antibody-discovery approaches, is a notable outlier for its contacts in the CXCR4 pocket [47]. We observe that, according to our predicted model, the backbone of antibody CXCR4_23 extends more than 3 Å farther into that pocket, highlighting the generative capability of the platform.

#### 2.6.3 Antibodies Designed to Agonize MRGPRX2

We obtained and validated an antibody hit against MRGPRX2, a key mediator of inflammatory and allergic mast-cell reactions. One natural ligand, cortistatin-14 (CST-14), activates MRGPRX2, triggering G protein-dependent calcium mobilization, mast cell degranulation and release of histamine, proteases, lipid mediators, and inflammatory cytokines [53–55]. According to the predicted model, the HCDR3 loop of the highest-affinity screening hit, MRGPRX2_15, engages the MRGPRX2-binding cavity via multiple CDR contacts (Fig. 6a). The dose-dependent binding of MRGPRX2_15 showed an EC_50_ of 84.5 nM, as determined by flow cytometry-based cell-surface measurements to overexpressing cells (Fig. 6b).

**Figure 6.**
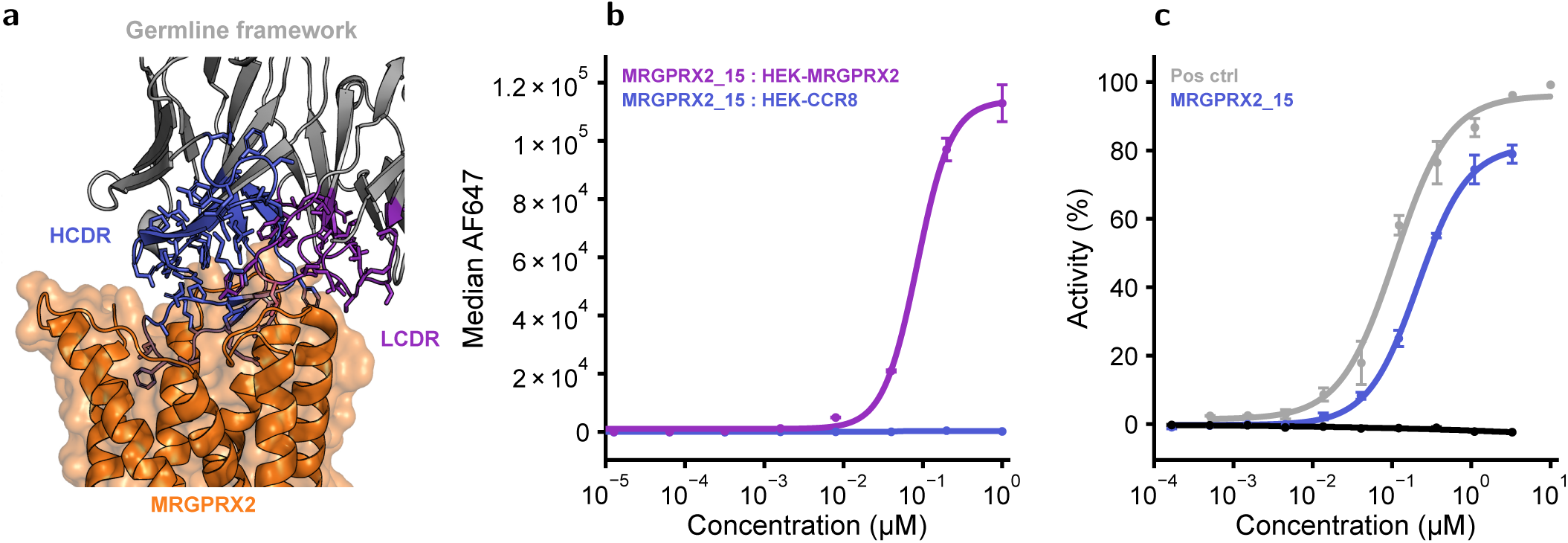
Characterization of designs to MRGPRX2. (a) Predicted binding pose of designed antibody MRGPRX2_15. (b) Concentration-dependent binding of MRGPRX2_15 to MRGPRX2-overexpressing cells (purple) measured by flow cytometry. The binding response was fit using nonlinear regression to determine the cell binding EC_50_ values. Binding of the same antibody sequence to CCR8-overexpressing cells (blue) was evaluated as an off-target cellular specificity control. (c) Functional agonism of MRGPRX2 signaling was assessed using the MRGPRX2 PathHunter® β-Arrestin recruitment assay. Percent activity was calculated relative to the maximum signal from the positive control. Curves were fit by nonlinear regression to determine functional potency. Negative control is represented as black; positive control is grey; and MRGPRX2_15 is blue.

The MRGPRX2 antibodies were designed to an active conformation of the receptor bound to ligand (PDB 7S8L [53]), with the aim of generating a full-length antibody that activates MRGPRX2. We tested the effect of MRGPRX2_15 in a dose-dependent reporter-based MRGPRX2 activation assay. The antibody induced concentration-dependent, saturable receptor activation, with an EC_50_ of 213.5 nM. This potency was comparable to that of CST-14, which had an EC_50_ of 109 nM in the same reporter assay (Fig. 6c). The CDR loops of MRGPRX2_15 are predicted to interact with the same receptor regions contacted by the natural ligand (Supplementary Fig. S12). The predicted binding pose and observed agonist activity are consistent with stabilization of the active MRGPRX2 conformation inducing downstream β-arrestin signaling. To our knowledge, this is the first report of a designed, full-length IgG antibody that activates a GPCR.

## 3 Discussion

This study demonstrates that our antibody design platform can produce full-length IgGs with high-affinity binding, functional activity, and favorable developability profiles across a broad range of therapeutic targets, including challenging GPCRs. The platform’s generative pipeline is powered by Pythia-1, an all-atom generative model that reasons jointly over amino acid sequence and three-dimensional atomic structure, under explicit design constraints. Design begins with requirements for both target recognition and key antibody properties, allowing functional objectives and therapeutic development considerations to guide candidate generation.

We show that antibodies can be designed for defined functional outcomes by specifying where they bind and the target conformation they recognize. Targeting the receptor-binding surfaces of IL11 and β-NGF yielded antagonists with single-digit nanomolar inhibitory potency in cell-based reporter assays, with β-NGF antagonism also confirmed in a cell-proliferation assay. For GPCRs, we selected receptor conformations according to the intended functional activity. Antibodies designed against an inactive CCR8 structure inhibited ligand-induced signaling, whereas an antibody designed against an active MRGPRX2 structure induced β-arrestin recruitment. Our approach overcomes longstanding barriers to designing functional, full-length antibodies against receptors with limited extracellular surface area and conformationally dynamic binding sites.

Key developability and manufacturability requirements were built into the initial design. We specified human germline frameworks, constrained CDR lengths to ranges observed in human antibodies, and guided amino acid sequence generation to limit known liabilities. The characterized designs compared favorably with clinical-stage and FDA-approved antibody controls across expression, thermal stability, surface hydrophobicity, and other biophysical measurements. These findings demonstrate that our technology can produce antibodies with key properties expected of therapeutic lead candidates from the outset, without requiring extensive downstream engineering to establish favorable biophysical profiles.

We demonstrate that our platform can produce high-affinity, functional antibodies in a single computational design round, with no more than 48 candidates experimentally screened per target. The computational workflow can generate candidates in a single day, and initial experimental screening was completed in less than three weeks. Generating functional antibodies without large experimental screens or iterative optimization substantially shortens the path to therapeutic lead candidates.

To our knowledge, this study provides the first demonstration of both GPCR agonism and antagonism by *de novo* designed full-length IgG antibodies. Together with functional antibodies against soluble proteins, this result establishes our platform’s ability to deliver full-length antibodies across diverse therapeutic targets, with functional activity extending from ligand neutralization to receptor inhibition and activation. *In vivo* assessment of pharmacokinetics, efficacy, and safety will be required to evaluate the suitability of these candidates for therapeutic development. We provide the complete sequences of all lead antibodies presented in this study to support independent validation and further research. By turning some of antibody discovery’s most demanding challenges into tractable design problems, Pythia-1 opens new opportunities to develop medicines previously beyond reach.

## 5 Materials and Methods

### 5.1 Target and epitope specifications

Structures were collected for the soluble and GPCR targets summarized in Fig. 1b. Binding epitope regions were derived from known ligand-target complexes in the PDB, experimental evidence reported in the literature, surface analysis or a combination of these. These definitions were subsequently refined manually to favor desirable binding modes and preserve critical structural features such as disulfide bonds. For some GPCRs, we carefully selected residues to facilitate deep binding into their orthosteric pockets. In addition, unresolved regions potentially contributing to the interface, as well as regions displaying apparent modeling artifacts, were refolded jointly with antibody generation.

### 5.2 Antibody specifications

The full-atom generative model was conditioned on frameworks derived from human germline antibody structures and predefined CDR boundaries. The CDRs were designed to form *de novo* interfaces with the target, without prescribed residue compositions, apart from the sequence-motif constraints described below. CDR lengths were jointly sampled from distributions derived from OAS [23] to provide combinations representative of the human antibody repertoire.

### 5.3 Generative pipeline

The generative pipeline integrates all-atom structure generation, sequence design, and candidate selection within a unified workflow depicted in Fig. 1a. It takes as input the three-dimensional structure of a target, a user-defined epitope, and an antibody scaffold, IgG in this study, to jointly generate candidate binder sequences and all-atom structures.

The first component of the pipeline is Pythia-1, a conditional all-atom flow-matching model [56] that jointly designs backbone geometry, sequence, and side-chain conformations. The model was trained in a multitask setting on diverse generative tasks across protein monomers, multimers, and antibody-antigen complexes. At inference, structures are sampled using the learned velocity field within the stochastic interpolants framework [57].

An independent confidence model was trained to predict the structural accuracy of generated designs [58–60]. Predicted scores include pLDDT, PAE and PDE. The pipeline iteratively refines and selects candidates via a beam search strategy.

Sequence can optionally be redesigned through backbone-conditioned sampling using a separate continuous-time Markov chain (CTMC) model [61]. CTMC sampling can be constrained by sequence motif rules to reduce liabilities such as N-linked glycosylation and deamidation-prone and isomerization motifs, among others. The final sequence pool can include both co-generated sequences from Pythia-1 and sequences generated by the CTMC model. Generated designs undergo a self-consistency check in which candidate sequences are refolded using Pythia-1 to assess recovery of the intended binding geometry. Final confidence metrics are evaluated on the refolded structures.

We applied a filtering strategy combining sequence-based, physics-based, and predicted confidence metrics to prioritize high-quality designs. Sequence-based filters screened for known sequence liabilities, including undesired disulfide-bonding patterns and glycosylation motifs, complementing the constraints applied during sequence design. Physics-based metrics assessed the geometric and energetic quality of the antibody-target interface, including steric clashes, electrostatic interactions, and other non-covalent interactions.

### 5.4 Dataset

Our generative models were trained on a dataset comprising structures available in the Protein Data Bank (PDB) as of June 2025 and high-confidence single-chain monomer predictions from AlphaFold2, totaling approximately 90 million entries. Proteins were clustered by sequence and structural similarity, with additional clustering by CDR geometry for SAbDab entries [29]. Training and test sets were constructed such that proteins sharing a cluster under any criterion were assigned to the same set. To minimize data leakage and support a fair evaluation, structures from the soluble panel, along with homologs belonging to their respective sequence or structural clusters, were removed from the training set. An analogous filtering step was not applied to the GPCR panel because most GPCR targets lacked antibody-bound structures in the PDB. However, a subsequent analysis confirmed that most GPCR structures specified as an input to Pythia-1 were not present in the training set, with the exception of CCR8 antagonist and C-X-C chemokine receptor type 7 (CXCR7) structures. Nevertheless, designs for these particular cases showed substantial sequence novelty (see Results).

### 5.5 Novelty and Diversity

The novelty and diversity of our designs were calculated within the subsets of all designs, primary hits, or verified hits for each given target.

### 5.6 Sequence and Structural Novelty

Design novelty was evaluated at sequence and structural levels following the method used by Chai 2 [11], using structures available in SAbDab. Sequence novelty was quantified as the minimum edit distance between each design’s concatenated CDRs and those of all references [62]. For structural novelty, we first identified reference complexes containing antigens similar to the target by sequence or structure. After superimposing target chains, we calculated the Cα RMSD of the antibody heavy-chain framework and reported the minimum value across the retrieved complexes. To evaluate structural diversity, we calculated sequence-aligned pairwise antibody RMSDs after superimposing target chains and performed average-linkage agglomerative clustering at a 3 Å cutoff [27]. Structural diversity is reported as the ratio of the number of clusters to the total number of designs in the subset.

### 5.7 Antibody Cloning and Expression

Synthetic genes encoding variable domains were ordered as codon-optimized DNA fragments and cloned into human IgG1 expression vectors. Antibodies were transiently expressed in ExpiCHO suspension systems under serum-free conditions. Heavy and light chain plasmids were transfected at equimolar ratios using PEI-based or proprietary transfection reagents. Cell cultures were maintained for 4–7 days depending on expression kinetics. Supernatants were clarified by centrifugation and 0.22-µm filtration prior to purification.

### 5.8 HTP-BLI Binding Assay Using CHO Supernatants

Constructs encoding full-length IgG1 antibody heavy and light chains were transiently transfected into 1-mL CHO cell cultures maintained in serum-free medium. Cells were cultured at 37 *^◦^*C and 5% CO_2_ with shaking, and antibodies were expressed and secreted into the culture supernatant for 3–5 days following transfection. Antibody binding was assessed directly from crude supernatants by biolayer interferometry (BLI) using an Octet RED384 instrument. Human IgG antibodies were immobilized onto AHC2 (Anti-Human IgG Fc Capture 2.0) or Protein A biosensors and subsequently exposed to monomeric antigen (with the exception of β-NGF which is a homodimer) at a single concentration of 1000 nM, to monitor association. Dissociation was monitored in the baseline buffer (PBS containing 0.03% Tween-20, pH 7.2). Measurements were performed at 30 *^◦^*C with shaking at 1,000 rpm. Data were processed using Octet Analysis Studio/OctetRED BLI Discovery (v12.2), and binding was evaluated based on the overall BLI response. Antibodies producing a total BLI response of greater than 0.06 nm at the end of the 150 s association phase were classified as primary screen hits.

### 5.9 Antibody Production and Purification

Recombinant antibodies were transiently expressed in CHO cells cultured under serum-free conditions (37 *^◦^*C, 5% CO_2_, with shaking). Supernatants were harvested 3–5 days after transfection, clarified by centrifugation and 0.22-µm filtration, and purified using Protein A magnetic beads (AmMag Protein A, GenScript) according to the manufacturer’s protocol. Antibodies were eluted under acidic conditions, neutralized, and buffer-exchanged into either 20 mM HEPES and 120 mM NaCl (pH 8.0) or 10 mM histidine and 120 mM NaCl (pH 6.0). Protein concentrations were determined by absorbance at 280 nm. Purity was assessed by nonreducing CE-SDS and SEC-HPLC. Endotoxin levels were *≤*0.1 EU/mg. Samples were aliquoted, flash frozen, and stored at *−*80 °C.

### 5.10 5-point BLI Affinity Determination

Binding of antibodies to respective antigens 14-3-3β (NBC1-18405, Novus), β-NGF (256GF100CF, R&D), CD19 (CD9-H52H2, Acro), IL11 (IL1-H5243, Acro), IL20 (13060-HNAE-100, Sino), IL7Rα-His (IL7-H52H7 Acro), and TrkA-His (11073-H08H-UE, Sino) was measured by BLI, with antigens reconstituted according to manufacturers’ instructions. Measurements were performed on an Octet RED384 or Octet R8 system (Sartorius) using ProA biosensors (Sartorius, Cat. No. 18-5010) in PBS containing 0.03% (v/v) Tween-20, pH 7.2. Antibodies (10 µg/mL) were captured to an immobilization level of 1.1–3.5 nm, followed by a baseline step in the assay buffer. Binding kinetics were measured at 30 *^◦^*C with shaking at 1,000 rpm using five 2-fold serial dilutions of antigen at the concentrations indicated in the figures. Association was monitored for 150 s and dissociation for 300 s in the assay buffer. Sensorgrams were analyzed using Octet Analysis Studio v12.2 (Sartorius) and globally fitted to a 1:1 Langmuir binding model to determine association and dissociation rates and equilibrium constants. Fit quality was assessed by *χ*^2^, R^2^, and visual inspection of the sensorgrams.

### 5.11 Surface Plasmon Resonance

Binding of monoclonal antibodies to programmed cell death protein 1 (PD-1) was measured by SPR on an Alto digital SPR system (Nicoya Lifesciences) using carboxymethyl-dextran (CMD) sensor cartridges (KC-CBX-CMD-16). An anti-human IgG Fc VHH capture reagent (ALTO-R-VHH-HR-KIT) was diluted to 2.5 µg/mL (167 nM) and covalently immobilized by standard EDC/NHS amine coupling. Purified antibodies were diluted up to 6-times molar excess ratio compared to the capture reagent in running buffer (HBS-T - 10 mM HEPES, 150 mM NaCl, 0.05% Tween-20, pH 7.4, supplemented with 5 mM EDTA and 0.1% BSA) and captured on the VHH surface to >300 RU. Recombinant human PD-1-His (PD1-H5221; ACROBiosystems) was injected at 500 nM, with association and dissociation monitored for 300 s and 500 s, respectively. All steps were performed at 25 *^◦^*C. Surfaces were regenerated between cycles with 10 mM glycine-HCl, pH 2.0. Pembrolizumab served as the positive control and HyHEL-10 (anti-hen egg lysozyme, IgG) as the non-binding control. Buffer-only injections were included for double referencing. Sensorgrams were double-referenced against the reference channel and buffer injections and analyzed in Nicoya analysis software. Binding was reported as response at the end of association normalized to capture level.

### 5.12 Differential Scanning Fluorimetry

Thermal stability was measured using the Protein Thermal Shift Dye Kit (Thermo Fisher Scientific, Cat. No. 4461146). Quadruplicate samples (2 µM) were prepared in 20-µL reactions containing 1x buffer and dye, loaded into optical PCR plates (Applied Biosystems, Cat. No. 4306737), and analyzed using a QuantStudio 5 system. Fluorescence (excitation, 470–490 nm; emission, 520–550 nm) was monitored during a 25–99 *^◦^*C thermal ramp. *T*_m_ values were determined by Boltzmann fitting and first-derivative analysis.

### 5.13 Dynamic light scattering (DLS)

Thermal aggregation and colloidal behavior were assessed on a Prometheus Panta system (NanoTemper Technologies) controlled by PR.Panta Control software. Before measurement, antibody samples were diluted to 1 mg/mL in respective low-fluorescence buffers (20 mM HEPES + 120 mM NaCl, pH 8.0 or 10 mM histidine, 120 mM NaCl, pH 6.0). 10 µL of each sample was loaded into Prometheus High Sensitivity capillaries (PR-C006) in triplicate, avoiding air bubbles and visible aggregates. Before the thermal ramp, an isothermal DLS measurement was acquired at 25 *^◦^*C to characterize the native-state size of each sample. DLS measurements were recorded using the integrated DLS module (405 nm laser, backscatter detection) with 10 acquisitions per data point. Autocorrelation functions were fitted by cumulant analysis to obtain the hydrodynamic radius (*R*_h_, nm) and polydispersity index (PDI), providing information on particle size distribution and colloidal homogeneity.

### 5.14 Hydrophobic Interaction Chromatography

HIC-HPLC separation was performed using a NanoChrom BioCore HIC-Butyl column. Antibodies were separated using a high-to-low salt gradient between mobile phases containing 1.5 mol/L ammonium sulfate, 50 mM sodium phosphate, and 5% isopropanol (pH 7.0) and 50 mM sodium phosphate and 20% isopropanol (pH 7.0). A 3-minute linear gradient was run at a flow rate of 1.8 mL/min with UV detection at 280 nm. Three micrograms of each antibody were injected. Pembrolizumab, lirilumab, tremelimumab, and adalimumab were included as retention-time controls.

### 5.15 Affinity-capture self-interaction nanoparticle spectroscopy (AC-SINS)

AC-SINS was performed as described previously [63]. Gold nanoparticles were coated with a polyclonal anti-human Fc capture antibody and blocked. Each antibody was tested at 700 nM in PBS, pH 7.2. Samples were incubated with the conjugated nanoparticles at room temperature before reading. Absorbance spectra were recorded on a plate reader at 2 nm intervals, and the wavelength of maximum absorbance (*λ*max) was identified for each well after subtraction of the blank (nanoparticles in buffer, no test antibody). Bococizumab served as a positive control (high self-association) and daclizumab as a negative control (low self-association) at a concentration of 700 nM, matching the test antibodies conditions.

### 5.16 Flow Cytometry Binding Assay Using CHO Supernatants and GPCR-Expressing Cells

Recombinant antibodies were produced in a CHO mammalian expression system. Constructs encoding antibody heavy and light chains were transiently transfected into 1 ml of CHO cells cultured in serum-free medium. Cells were maintained at 37 *^◦^*C and 5% CO_2_ with shaking and allowed to express and secrete antibodies for 3–5 days after transfection. Supernatants were then frozen.

Binding of antibody-containing supernatants to cell-surface GPCRs was assessed by flow cytometry using stable GPCR-overexpressing cell lines. Cells were harvested and distributed into U-bottom 96-well plates at 1 *×* 10^5^ cells per well. Cells were washed with 200 µL of phosphate-buffered saline (PBS) per well and incubated with 100 µL of CHO supernatant for 30 min at 4 *^◦^*C to minimize receptor internalization. Cells were then washed with 200 µL of cell-staining buffer to remove unbound antibody. Cell-associated human IgG was detected using an Alexa Fluor 647-conjugated anti-human IgG secondary antibody diluted to its working concentration in cell-staining buffer. Propidium iodide (PI; 1:2,000) was included in the secondary-antibody solution to permit exclusion of nonviable cells. Cells were incubated with 100 µL of the secondary-antibody mixture for 30 min at 4 *^◦^*C protected from light, washed, resuspended in 100 µL of cell-staining buffer, and analyzed using an Attune flow cytometer (Thermo Fisher Scientific).

Antibody binding was quantified as the median fluorescence intensity (MdFI) in the Alexa Fluor 647 detection channel. In primary binding screens, hits were identified on a plate-by-plate basis using a robust z-score. The median Alexa Fluor 647 MdFI of the antibody population on each plate was used as the measure of central tendency, and variability was estimated using the median absolute deviation (MAD). The robust standard deviation was calculated as robust SD = 1.4826 *×* MAD, where MAD = median (|xi - median(x)|). The robust z-score for each antibody was then calculated as robust Z = (xi - plate median)/(robust SD), where xi represents the Alexa Fluor 647 MdFI of the test antibody. Antibodies producing an Alexa Fluor 647 MdFI more than 10 robust standard deviations above the plate median, corresponding to a robust z-score >10, were classified as primary binding hits.

### 5.17 Concentration-Response Binding and Cell-Surface Specificity Assessment

Antibodies identified as hits in the primary flow cytometry screen were purified and characterized further for concentration-dependent binding to the corresponding GPCR-expressing cell line. Purified antibodies were tested in a 10-concentration, fivefold serial dilution from 1,000 to 0.000512 nM. Cells were plated at 1 *×* 10^5^ cells per well in U-bottom 96-well plates and stained using the flow cytometry procedure described above. Briefly, cells were incubated with test antibodies for 30 min at 4 *^◦^*C and washed, and bound antibodies were detected using an Alexa Fluor 647-conjugated anti-human IgG secondary antibody. Propidium iodide was included during secondary staining to permit exclusion of nonviable cells. Cells were analyzed using an Attune flow cytometer (Thermo Fisher Scientific), and antibody binding was quantified as MdFI in the Alexa Fluor 647 detection channel. To assess cell-surface binding specificity, each antibody was tested in parallel across the same concentration range against a control cell line derived from the same parental background but engineered to overexpress an unrelated GPCR. Matched parental cell backgrounds enabled assessment of target-dependent binding while controlling for nonspecific interactions with the host cell line, cell-surface components, or features associated with GPCR overexpression.

Concentration-response curves were generated from Alexa Fluor 647 MdFI values. When concentration-dependent binding was observed, curves were fitted by nonlinear regression using a four-parameter logistic model to estimate the apparent half-maximal binding concentration (apparent EC_50_) and maximal binding response. The apparent EC_50_ is the antibody concentration producing 50% of the fitted maximal cell-associated fluorescence under the assay conditions and was used as a comparative measure of cell-surface binding potency rather than a direct measure of equilibrium binding affinity. Antibodies were considered target-selective when concentration-dependent binding was observed on the intended GPCR-expressing cell line and reduced or negligible binding was observed on the matched off-target GPCR-expressing control cell line.

### 5.18 HEK293-F Transfection and Binding

HEK293-F cells were subcultured 24 h before transfection to ensure logarithmic growth, and targets were tagged with eGFP to monitor expression. On the day of transfection, cells were diluted in FreeStyle HEK293 Expression Medium to 1*×*10^6^ cells/mL at >95% viability. Depending on the target, 1 or 2 µg of plasmid DNA per 10^6^ HEK293-F cells was diluted in Opti-MEM. The 293fectin reagent was prepared separately in Opti-MEM at a 2:1 reagent-to-DNA ratio. Diluted transfection reagent was added gently to the diluted DNA, and the mixture was incubated for 20 min at room temperature to allow complexes to form. The DNA-293fectin complex was then added dropwise to the cell suspension. Cells were incubated for 48 h at 37 *^◦^*C and 8% CO_2_ on an orbital shaker at 125 rpm. Cells that successfully expressed the target were eGFP-positive.

Transfected cells were plated in U-bottom 96-well plates at 1–2*×*10^5^ cells per well. Cells were washed with 200 µL of PBS per well and incubated with 100 µL of 1x titrated antibody for 30 min at 4 *^◦^*C. Cells were washed with 200 µL of cell-staining buffer, and bound antibody was detected using 100 µL of 1x anti-human IgG secondary antibody diluted in cell-staining buffer containing PI (1:2,000). After incubation for 30 min at 4 *^◦^*C, cells were washed, resuspended in 100 µL of cell-staining buffer, and analyzed using an Attune flow cytometer.

### 5.19 TF-1 Cell-Proliferation Assay

The functional activities of purified antibody hits were evaluated using a β-NGF-dependent TF-1 cell-proliferation assay. TF-1 cells were maintained in RPMI supplemented with 10% fetal bovine serum (FBS) and 2 ng/mL granulocyte-macrophage colony-stimulating factor. On the day before the assay, cells were harvested, counted, washed once with RPMI, and cells cultured overnight in reduced serum medium (RPMI, 5% FBS) to reduce basal proliferation.

On the day of the assay, serum-starved cells were harvested, counted, and resuspended in RPMI containing 5% FBS. Cells were seeded in U-bottom 96-well plates at 1*×*10^4^ cells per well in 90 µL and incubated for 60 min at 37 *^◦^*C and 5% CO_2_ before treatment. Purified test antibodies and recombinant human β-NGF (rh β-NGF) were prepared separately at 20x their intended final assay concentrations in RPMI containing 5% FBS. Equal volumes of the 20x antibody and 20x rh β-NGF solutions were combined to generate a 10x treatment mixture and preincubated for 30 min at room temperature. A 10-µL aliquot of the antibody/β-NGF mixture was added to each well, resulting in a final volume of 100 µL. Cells were incubated for 40 h at 37 *^◦^*C and 5% CO_2_.

Cell proliferation and viability were quantified using CellTiter-Glo 2.0 (Promega), which measures cellular ATP as a luminescent indicator of metabolically active cells. After the 40-h treatment, assay plates were equilibrated to room temperature for approximately 30 min. CellTiter-Glo 2.0 reagent was equilibrated separately to room temperature protected from light and added at 100 µL per well. Plates were mixed for 2–3 min to induce cell lysis and incubated for an additional 10 min at room temperature in the dark. A 100-µL aliquot from each well was transferred to a solid white 96-well plate, and luminescence was measured using a Varioskan multimode plate reader (Thermo Fisher Scientific). The effect of each antibody on β-NGF-dependent TF-1 proliferation was determined by comparison of luminescence values with appropriate β-NGF-stimulated and unstimulated control wells.

### 5.20 TrkA Cell-Based Kinase β-NGF Neutralization Assay

The ability of purified anti-β-NGF antibody hits to inhibit β-NGF-mediated TrkA activation was evaluated using the LeadHunter TRKA Human RTK Kinase Cell-Based Assay (Eurofins DiscoverX, Fremont, CA, USA) in antagonist (anti-ligand) format. The PathHunter functional receptor tyrosine kinase assay uses enzyme-fragment complementation to measure ligand-dependent receptor activation. Full-length TrkA is expressed with the ProLink (PK) fragment of β-galactosidase fused to its intracellular C-terminal region and an SH2 domain-containing protein fused to the complementary Enzyme Acceptor (EA) fragment. Ligand-induced TrkA activation and phosphorylation recruit the SH2-EA fusion protein to the receptor, resulting in complementation of the β-galactosidase fragments and generation of a chemiluminescent signal.

PathHunter cells were expanded from frozen stocks according to standard procedures and seeded in white-walled 384-well microplates in a volume of 20 µL per well. Test antibodies were prepared as 6*×* solutions in assay buffer and combined with recombinant human β-NGF at its predetermined EC_80_ concentration. Antibody and β-NGF were preincubated for 60 min at room temperature prior to addition to cells. Following preincubation, 10 µL of the resulting 3*×* antibody/β-NGF mixture was added to each well, and cells were incubated for 90–180 min at 37 *^◦^*C or room temperature according to the assay-specific conditions. The β-NGF EC_80_ concentration used for the TrkA assay was 0.016 µg/mL.

Following stimulation, 15 µL of PathHunter Detection Reagent cocktail was added to each well, and plates were incubated for 1 h at room temperature. Chemiluminescence was measured using a PerkinElmer EnVision™ multimode plate reader. Antibody activity was analyzed using the CBIS data analysis suite (ChemInnovation, CA, USA). Percentage inhibition of β-NGF-induced TrkA activation was calculated relative to vehicle and EC_80_ β-NGF controls according to:

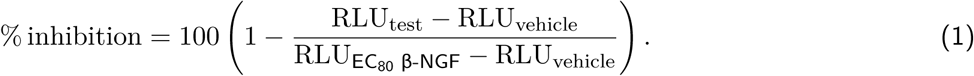

Concentration-response curves were generated from normalized inhibition values, and IC_50_ values were determined where concentration-dependent inhibition was observed.

### 5.21 HepG2 IL11 Reporter Assay

The antagonistic activities of purified anti-IL11 antibodies were evaluated using an IL11-responsive HepG2 reporter cell assay (Eurofins DiscoverX). Cells were maintained under standard culture conditions and used at approximately 70–80% confluence during logarithmic growth. On the day of assay setup, cells were washed with calcium- and magnesium-free PBS and dissociated using AssayComplete Cell Detachment Reagent. Detached cells were collected in culture medium, counted, and centrifuged at 300 *×* g for 5 min. The cell pellet was resuspended in AssayComplete Cell Plating Reagent at 5.5*×*10^4^ cells/mL.

Cells were seeded in white, flat-bottom, tissue culture-treated 96-well plates at 5,000 cells per well in 90 µL and incubated for 4 h at 37 *^◦^*C and 5% CO_2_ before treatment. Recombinant human IL11 (rhIL11) and test antibodies were diluted in AssayComplete Protein Dilution Buffer. The rhIL11 protein was prepared at 20x the predetermined EC_80_ concentration, and antibodies were prepared as 20x solutions using an eight-point, 5-fold serial dilution series. Antibody titrations were tested in triplicate. Equal volumes of the 20x rhIL11 and 20x antibody solutions were combined and preincubated for 30 min at room temperature, generating a 10x treatment mixture. A 10-µL aliquot of each rhIL11/antibody mixture was subsequently added to the appropriate wells, resulting in a final assay volume of 100 µL per well. Plates were incubated for 16 h at 37 *^◦^*C and 5% CO_2_.

Following treatment, reporter activity was quantified by luminescence. A 1:1 mixture of Enzyme Acceptor (EA) reagent and lysis buffer was added at 20 µL per well, and plates were incubated for 15 min at room temperature. Substrate reagent was then added at 40 µL per well, followed by incubation for 1 h at room temperature protected from light. Luminescence was measured using a multimode luminescence plate reader with an integration time of 0.1–1 s per well.

Antibody-mediated inhibition of IL11 signaling was determined relative to rhIL11-stimulated and unstimulated control wells. Concentration-response curves were generated from normalized luminescence values and fit by nonlinear regression to determine the half-maximal inhibitory concentration (IC_50_), where applicable.

### 5.22 PathHunter β-Arrestin Recruitment Assays

GPCR-mediated β-arrestin recruitment was evaluated using PathHunter® β-Arrestin assays (Eurofins DiscoverX, Fremont, CA, USA). Assays were performed for CXCR4 and CCR8 in antagonist mode, MRGPRX2 in agonist mode, and CXCR7 in both agonist and antagonist modes.

The PathHunter assay uses enzyme fragment complementation to quantify recruitment of β-arrestin to activated GPCRs. Briefly, the target GPCR is fused to the ProLink™ (PK) fragment of β-galactosidase, while β-arrestin is fused to the complementary Enzyme Acceptor (EA) fragment. GPCR activation and subsequent β-arrestin recruitment bring the PK and EA fragments into proximity, restoring β-galactosidase activity. Enzyme activity is detected using a chemiluminescent substrate, providing a quantitative measure of receptor-dependent β-arrestin recruitment.

PathHunter cells expressing the indicated receptor were expanded according to the manufacturer’s standard procedures and seeded in white-walled 384-well microplates at a volume of 20 µL per well. For agonist-mode assays (MRGPRX2 and CXCR7), test antibodies were prepared as 5x intermediate dilutions in the assay buffer, and 5 µL was added to cells. Cells were incubated with test antibodies for at least 120 min at 37 *^◦^*C or room temperature, as specified for the individual assay. Agonist activity was normalized to the response produced by the assay-specific reference agonist, with vehicle-treated cells defining the minimum response.

For antagonist-mode assays (CXCR4, CCR8, and CXCR7), cells were incubated with serial dilutions of test antibodies using the standard PathHunter antagonist assay format and subsequently stimulated with an assay-specific agonist at a concentration selected to produce a robust β-arrestin recruitment response. For the CCR8 assay, CCL1 was used at its EC_80_ concentration. Antagonist activity was determined from the reduction in agonist-induced β-arrestin recruitment relative to agonist and vehicle controls.

Following completion of the receptor stimulation period, 15 µL of PathHunter Detection Reagent cocktail was added to each well of the final detection mixture. Plates were incubated for 1 h at room temperature and chemiluminescence was measured using a PerkinElmer EnVision™ multimode plate reader. Data were analyzed using the CBIS data analysis suite (ChemInnovation, CA, USA). For agonist-mode assays, percentage activity was calculated relative to the vehicle and maximal reference-agonist responses according to:

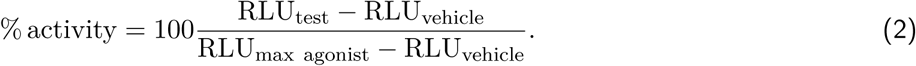

For antagonist-mode assays, responses were normalized to the agonist-stimulated and vehicle/background controls and expressed as percentage inhibition of agonist-induced β-arrestin recruitment. Concentration-response curves were generated to determine EC_50_ values for agonist activity or IC_50_ values for antagonist activity, where applicable.

## Data availability

The sequences of the antibodies with confirmed binding described in this study are available for download from the Pythia Labs website (https://pythialabs.com).

## Supplementary information

**Figure S1.**
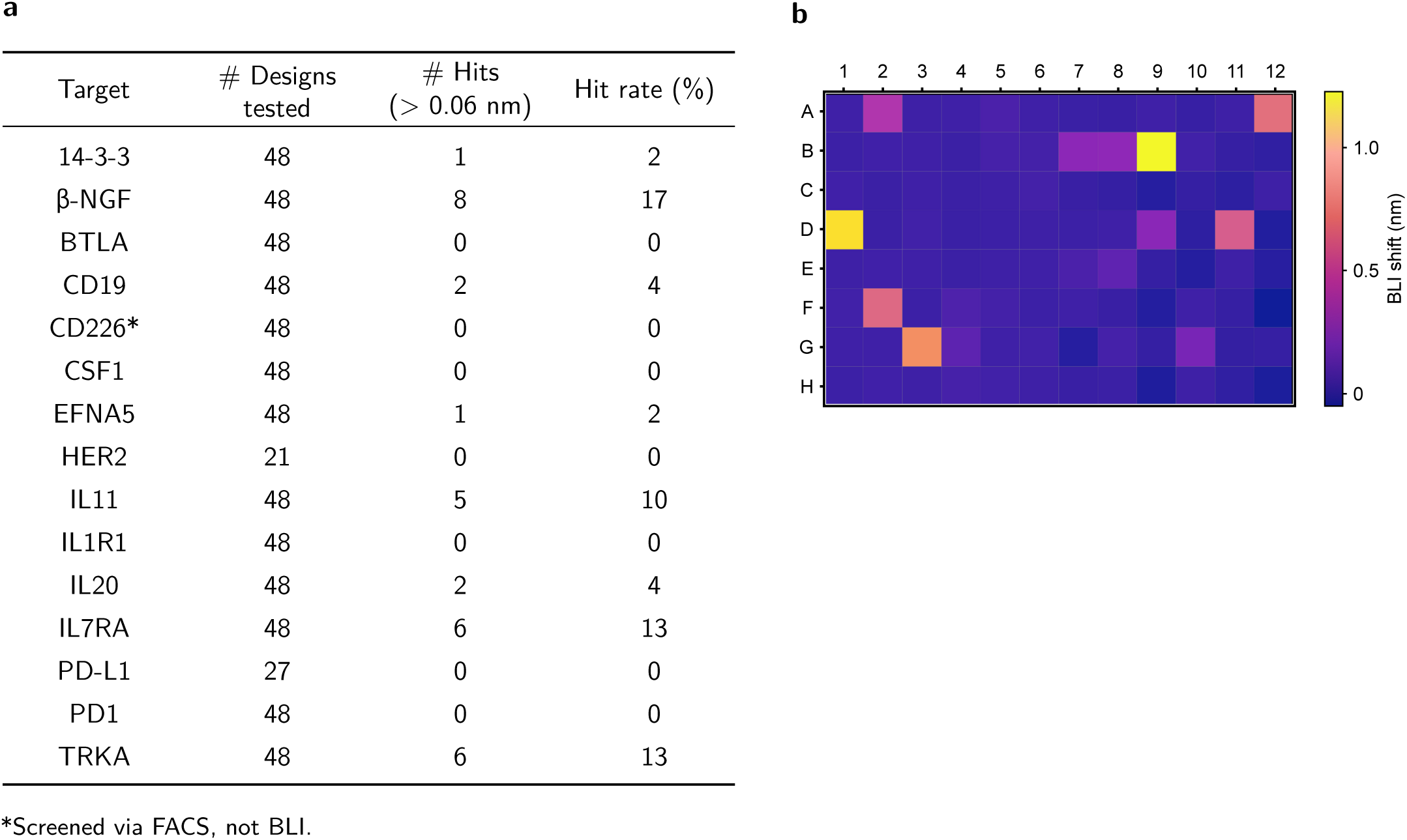
Primary screening hits and affinities. (a) Summary of soluble domain targets screened via HTP-BLI and resulting hits. (b) Example heatmap of BLI HTP screening data for soluble domain antigens (in this case, IL11 and β-NGF). Each square represents the total BLI response in nm at 150 seconds post antigen association to one antibody, with BLI response indicated by the color bar on the right side of the heat map. Antibodies with a signal above 0.06 nm were classified as a hit for follow-up studies.

**Figure S2.**
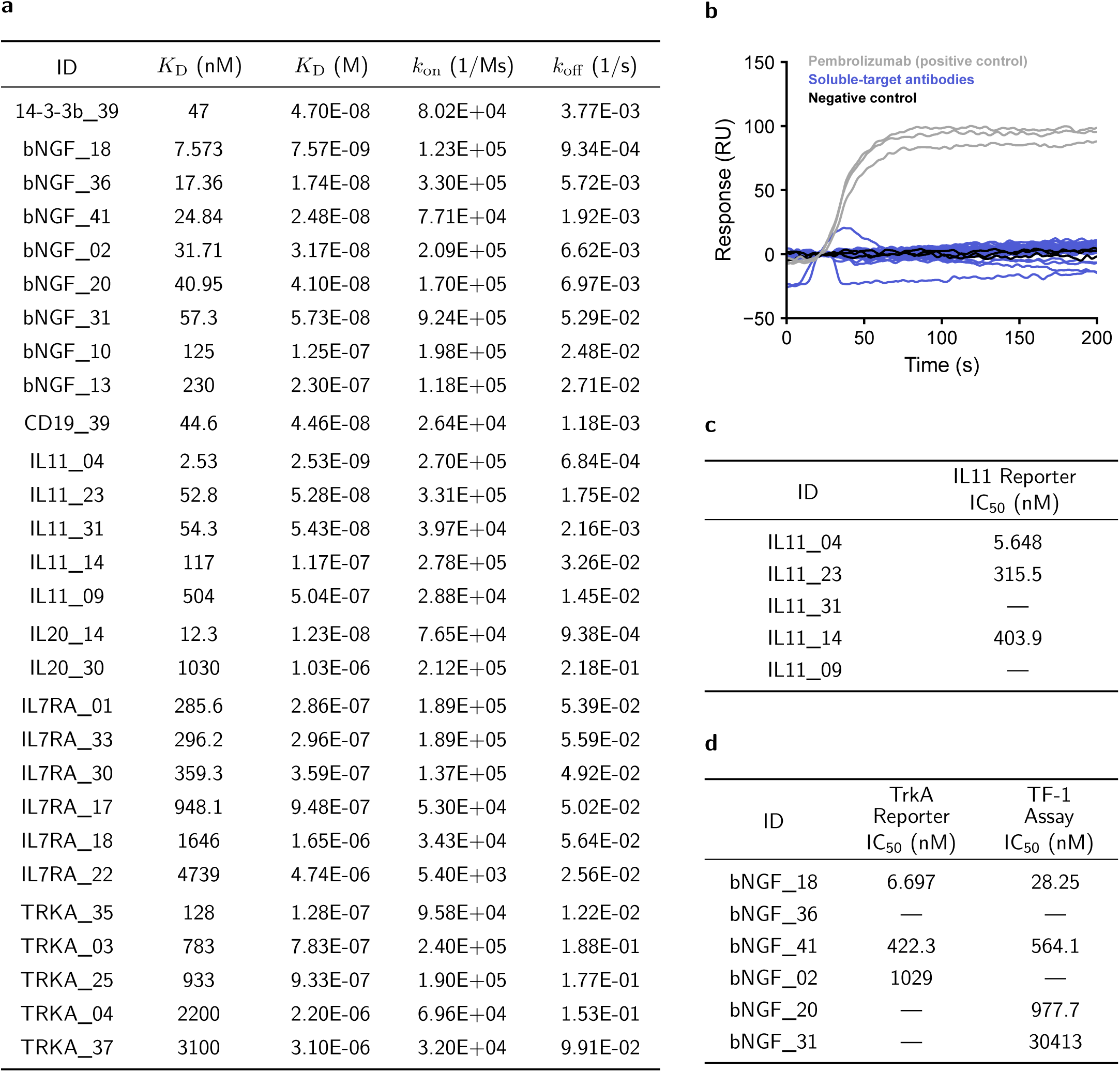
***De novo*** design of IgGs against soluble target domains yields hits with functional activity. (a) Summary of measured on/off rates and corresponding affinities for measured antibodies against their target antigens (β-NGF is a homodimer). Designs exhibiting no/very weak binding (> 5 uM) are not included. (b) SPR sensorgrams showing no detectable binding of soluble-target antibodies (blue) to the negative-control antigen PD1, compared with pembrolizumab as a positive control (grey) and a non-binding antibody as a negative control (black). For clarity, only the first 200 s of association are shown. (c) IC_50_ values of Pythia designed anti-IL11 antibodies using the PathHunter® Human IL11 (IL11RA) Signaling STAT3 Reporter Stable Cell Line Assay (HepG2) in the presence of IL11 at a concentration corresponding to EC_80_. (d) IC_50_ values of Pythia designed anti-β-NGF antibodies using the LeadHunter® TRKA Human RTK Kinase Cell Based Antagonist Assay in the presence of β-NGF at a concentration corresponding to EC_80_. Percent inhibition values were calculated relative to a β-NGF control alone. The same antibodies were also evaluated in a β-NGF-dependent TF-1 proliferation assay with dose responses measured by Cell Titer Glo 2.0 after 40 hr incubation.

**Figure S3.**
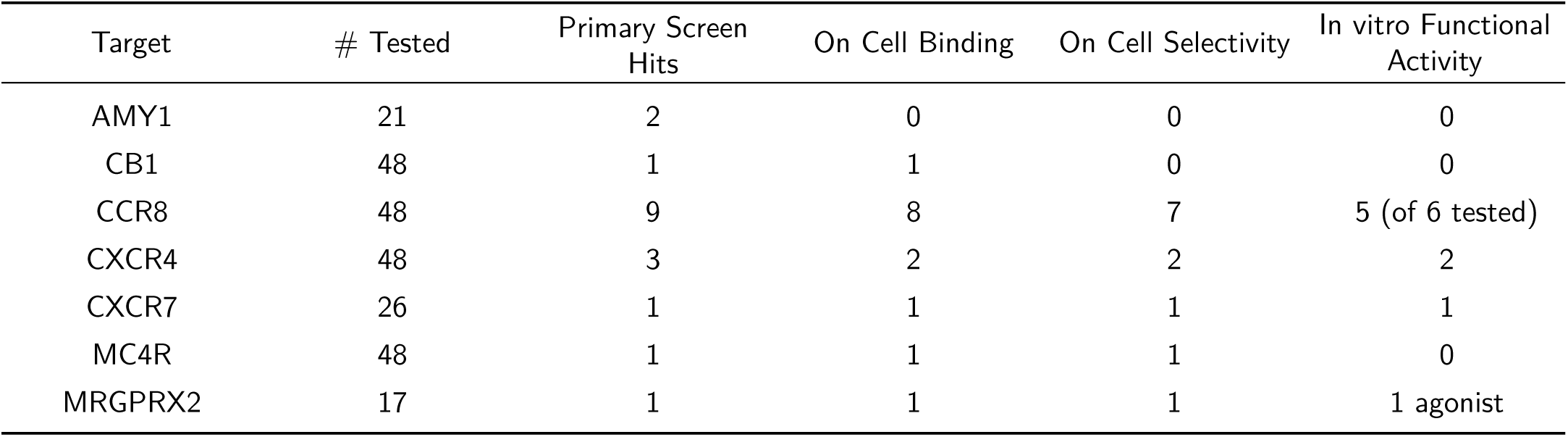
***De novo*** design of IgGs against GPCRs yields hits with functional activity. Summary of antibody progression through the validation cascade for each GPCR target. Columns indicate the number of sequences tested in the primary screen, primary screening hits, sequences demonstrating concentration-dependent cell binding, sequences meeting on-cell selectivity criteria, and sequences demonstrating activity in the corresponding *in vitro* functional assay. Parenthetical values indicate the number of sequences evaluated when not all candidates were tested at a given stage.

**Figure S4.**
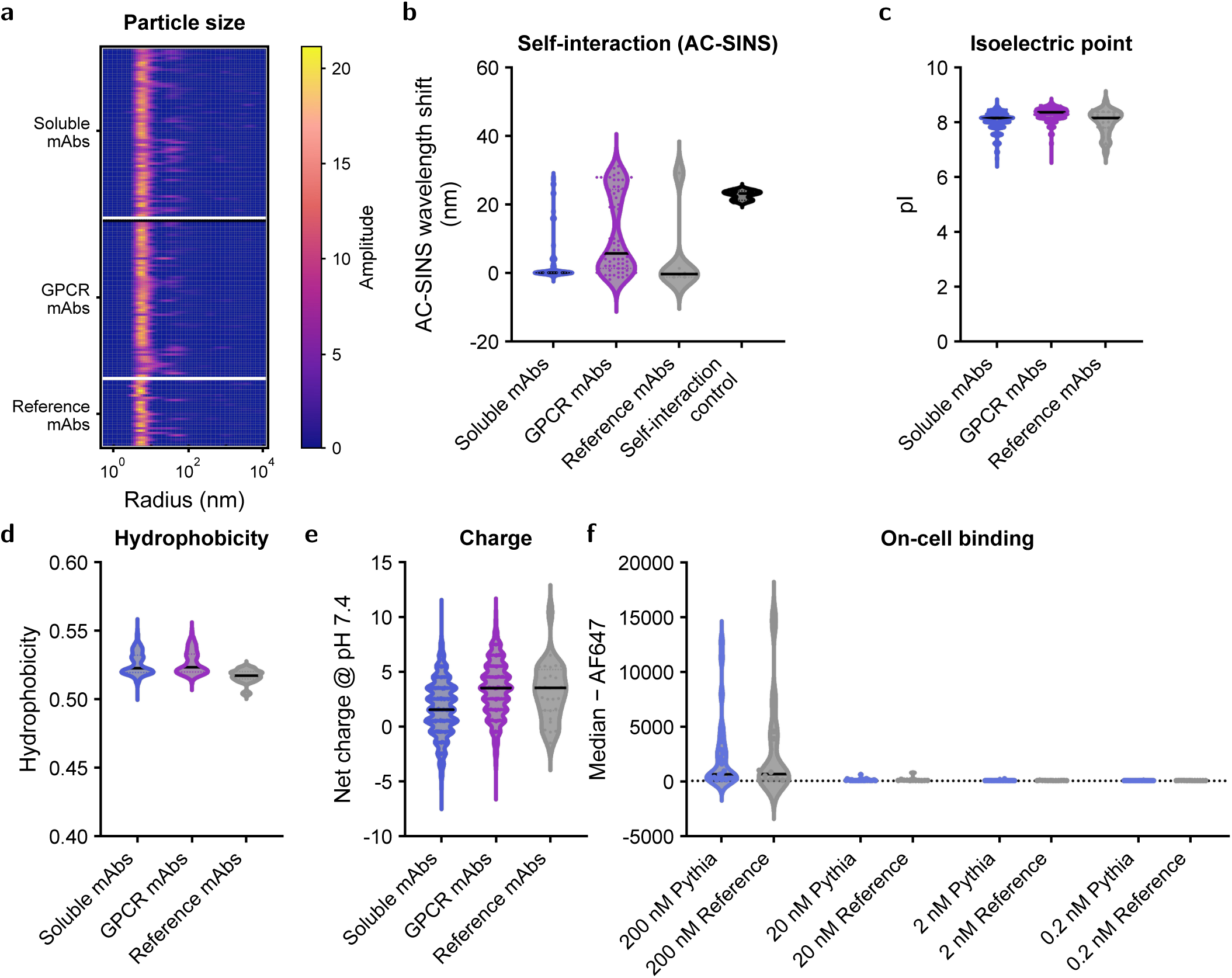
Developability assessment of Pythia mAbs compared with reference antibody sets. (a) Heat map showing particle size distributions for Pythia mAbs and reference mAbs as measured by dynamic light scattering. Scattering amplitude is plotted across particle radius, with warmer colors indicating higher abundance. A predominant signal at low nanometer-scale radius is consistent with monomeric antibody species, while larger-radius signal indicates higher-order particles or aggregates. (b) AC-SINS analysis showing self-interaction as measured by wavelength shift. (c–e) Predicted isoelectric point (pI) (c), hydrophobicity (d), and net charge at pH 7.4 (e) for Pythia designed antibodies relative to the reference antibody set. (f) Assessment of non-specific cell binding. Antibodies with confirmed binding to β-NGF, CD19, IL11, IL20, IL7RA, and TrkA were assessed for nonspecific binding to 293-F cells by flow cytometry at the indicated concentrations (200, 20, 2, and 0.2 nM). Pythia-designed antibodies (blue) were compared with reference antibodies (grey). Binding was quantified by median AF647 fluorescence intensity. Violin plots show the distribution across each antibody set, with black horizontal lines indicating group medians.

**Figure S5.**
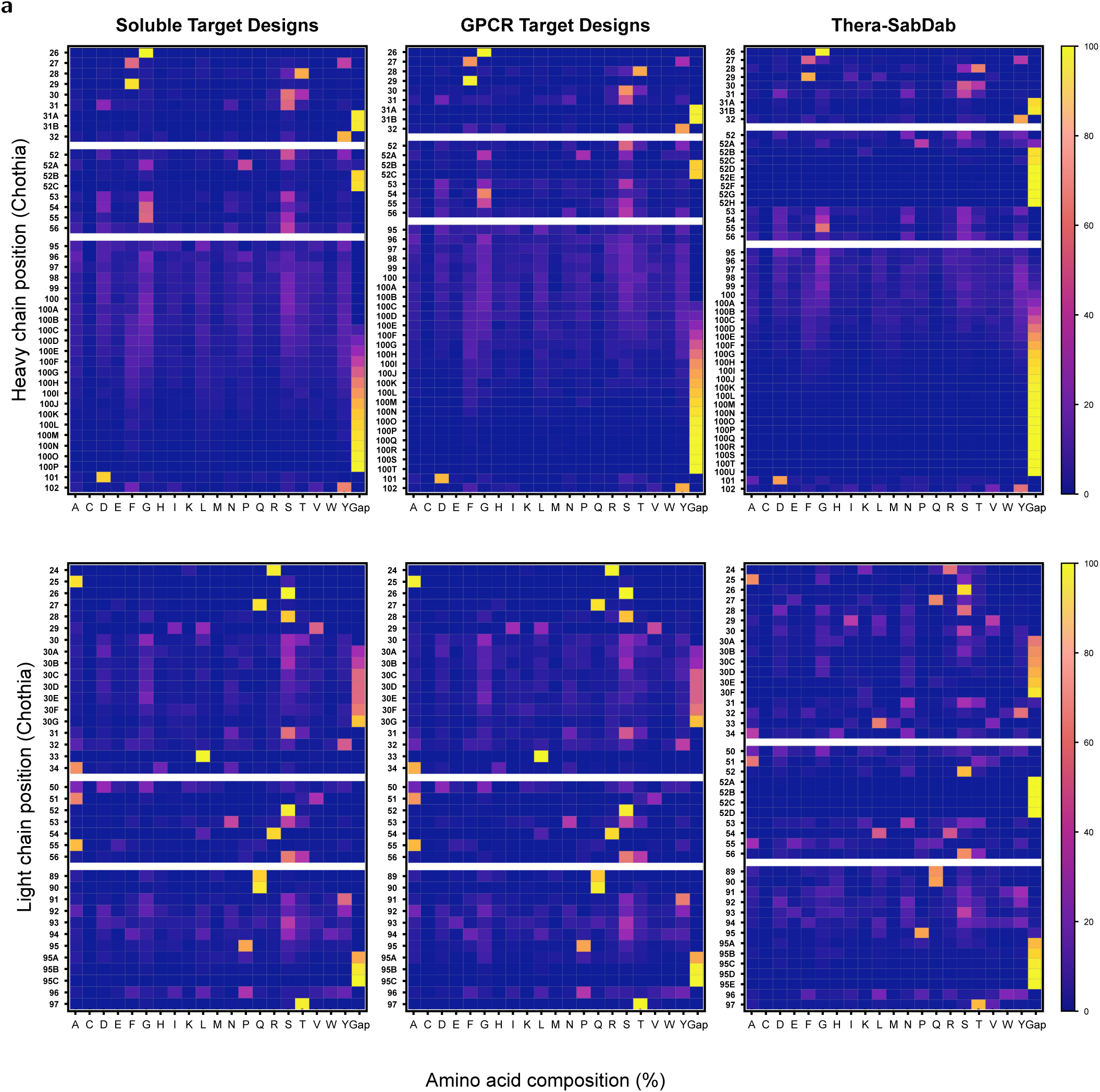
Amino acid composition of heavy and light chain CDRs. (a) Amino acid composition of antibody CDRs across Pythia-designed antibodies targeting soluble targets (left) and GPCR targets (middle) relative to Thera-SabDab (right), for both heavy chain (top) and light chain (bottom). Heat maps show the relative frequency of each amino acid at each position, presented as a percentage of total at each position. Warmer colors (yellow) indicate increased amino acid frequency, while cooler colors (blue) indicate decreased frequency. Overall patterns highlight similarities and differences in CDR composition associated with target class and provide a comparison to clinically validated therapeutic antibody repertoires.

**Figure S6.**
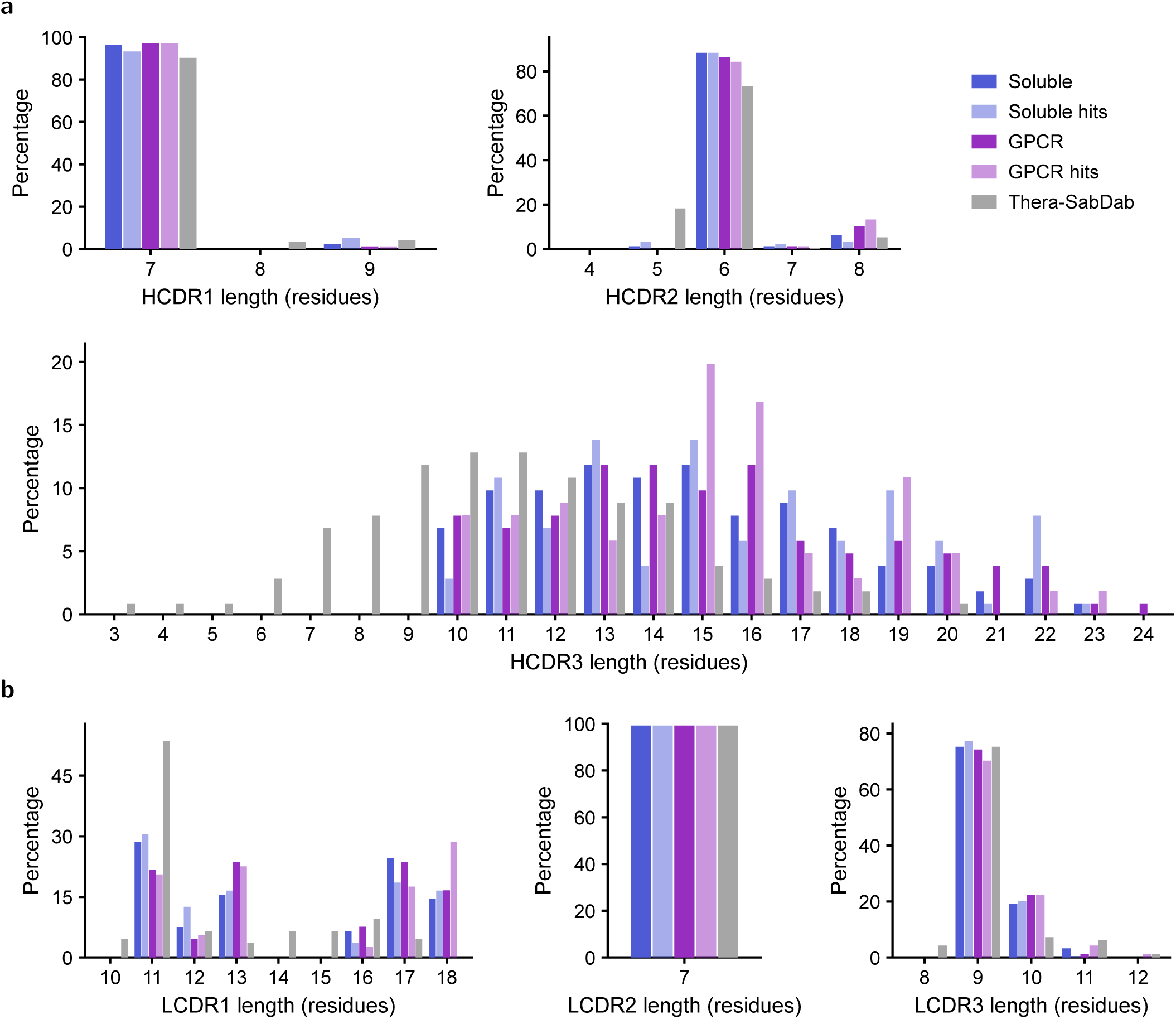
Amino acid CDR length distributions across antibodies targeting soluble protein domains and GPCRs compared with therapeutic antibodies from Thera-SabDab. **(a)** Heavy-chain CDR length distributions for HCDR1, HCDR2, and HCDR3 across soluble-target antibodies (dark blue), soluble-target hits (light blue), GPCR-target antibodies (dark purple), GPCR-target hits (light purple), and Thera-SabDab antibodies (grey). **(b)** Light-chain CDR length distributions for LCDR1, LCDR2, and LCDR3 across the same antibody groups. Bars indicate the percentage of antibodies within each dataset having the indicated CDR length, as defined by Chothia residue numbering.

**Figure S7.**
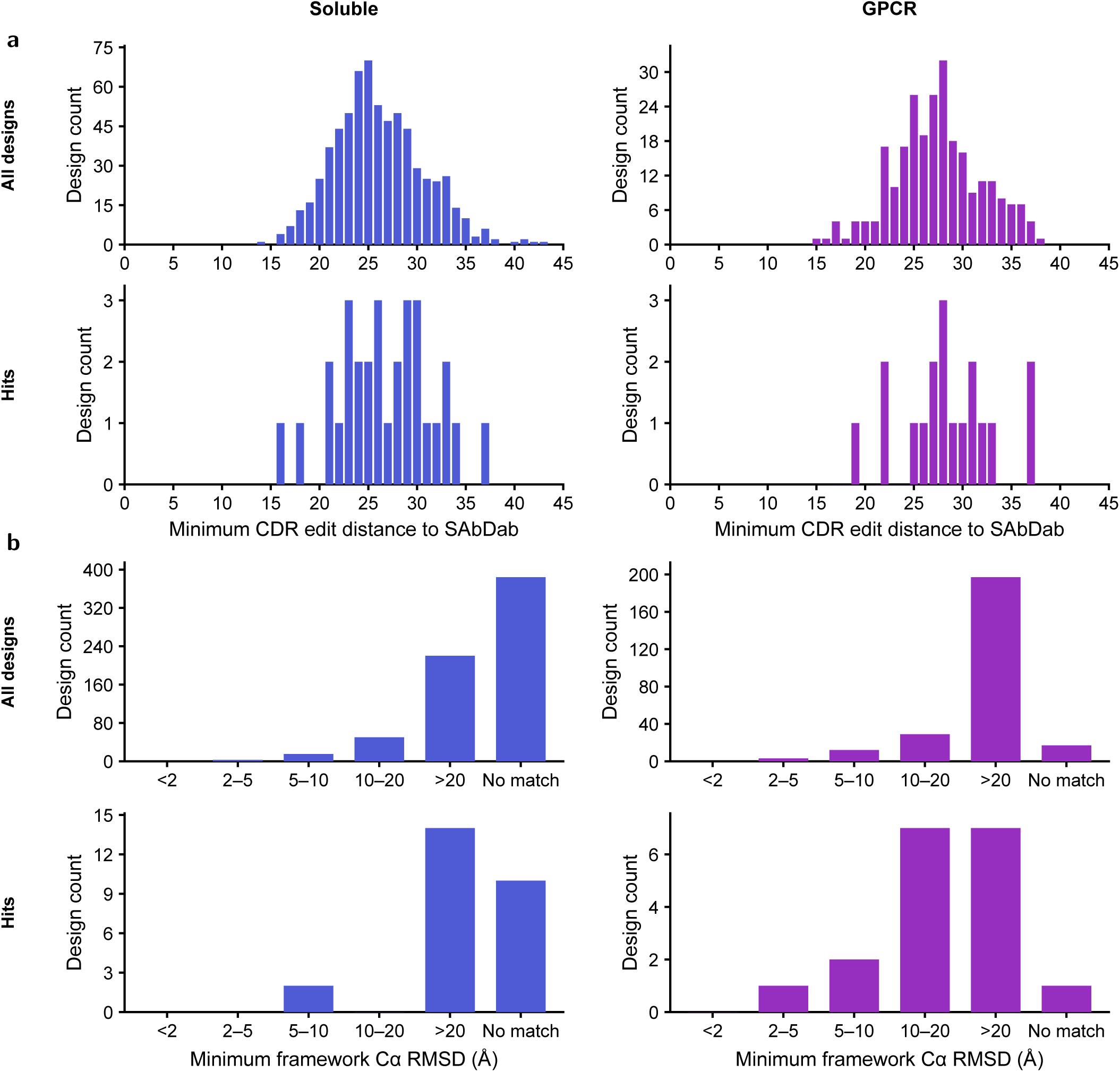
Sequence and structural novelty of Pythia-1 designed mAbs. (a) Sequence novelty of designed antibodies relative to known antibody sequences. The figure compares designs to external antibody references; within-set structural diversity is shown in Supplementary Fig. S8. Histograms show the distribution of the minimum CDR edit distance to SAbDab for antibody designs targeting soluble proteins (blue, left) and GPCRs (purple, right). Distributions are shown for all designs (top) and experimentally identified hits (bottom). Higher minimum CDR edit distances indicate greater sequence divergence from the closest SAbDab antibody CDR sequence. (b) Structural novelty of designed antibodies relative to known antibody structures. Bar plots show the distribution of minimum antigen-aligned framework Cα RMSD to SAbDab structures for antibody designs targeting soluble proteins (blue, left) and GPCRs (purple, right). Distributions are shown for all designs (top) and experimentally identified hits (bottom). Higher RMSD values indicate greater structural divergence from the closest antigen-aligned SAbDab antibody framework.

**Figure S8.**
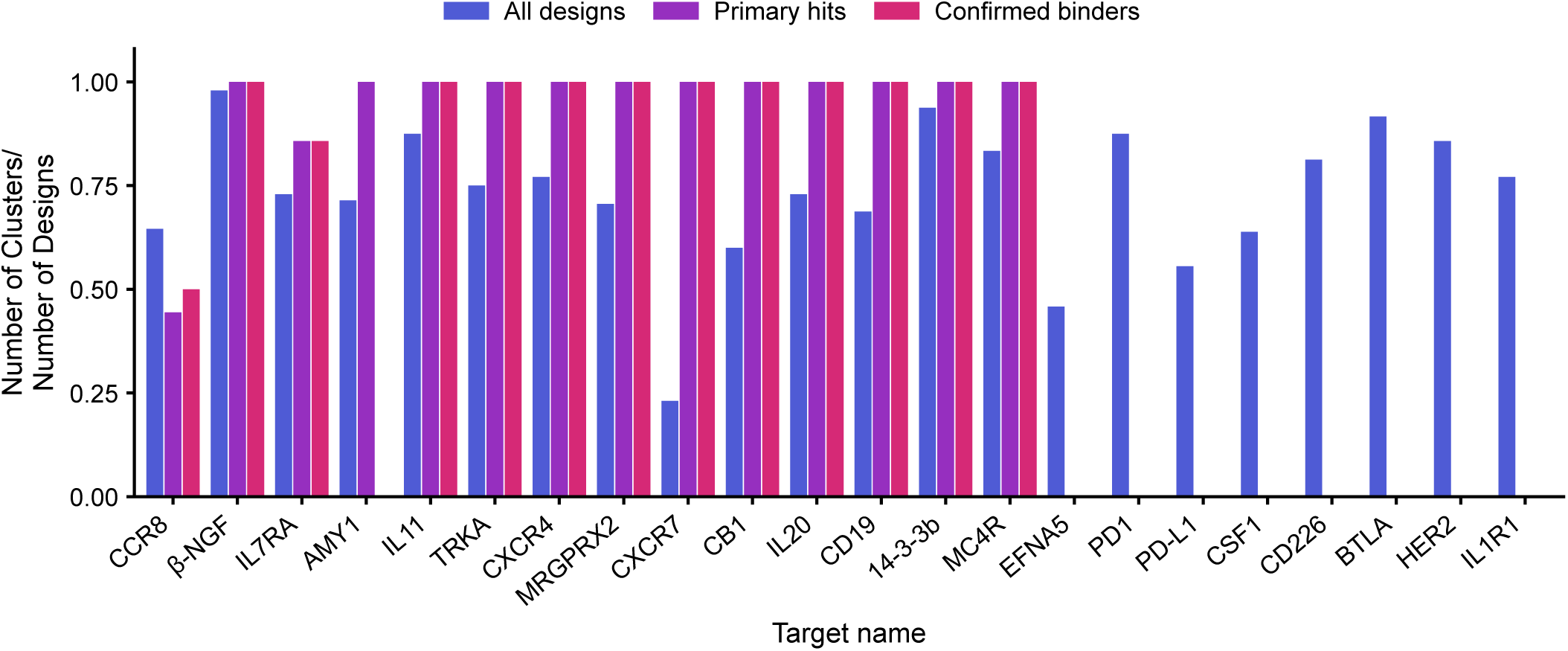
Diversity clustering of tested designs. Agglomerative clustering at 3 Å was performed for each subset of antibodies designed against each target. For each subset, including all computational designs, primary screening hits only, or confirmed binding hits only, sequence aligned pairwise RMSD were calculated for the antibody heavy and light chains including the CDR regions. This captures both pose and local CDR binding interactions in the diversity of the overall set.

**Figure S9.**
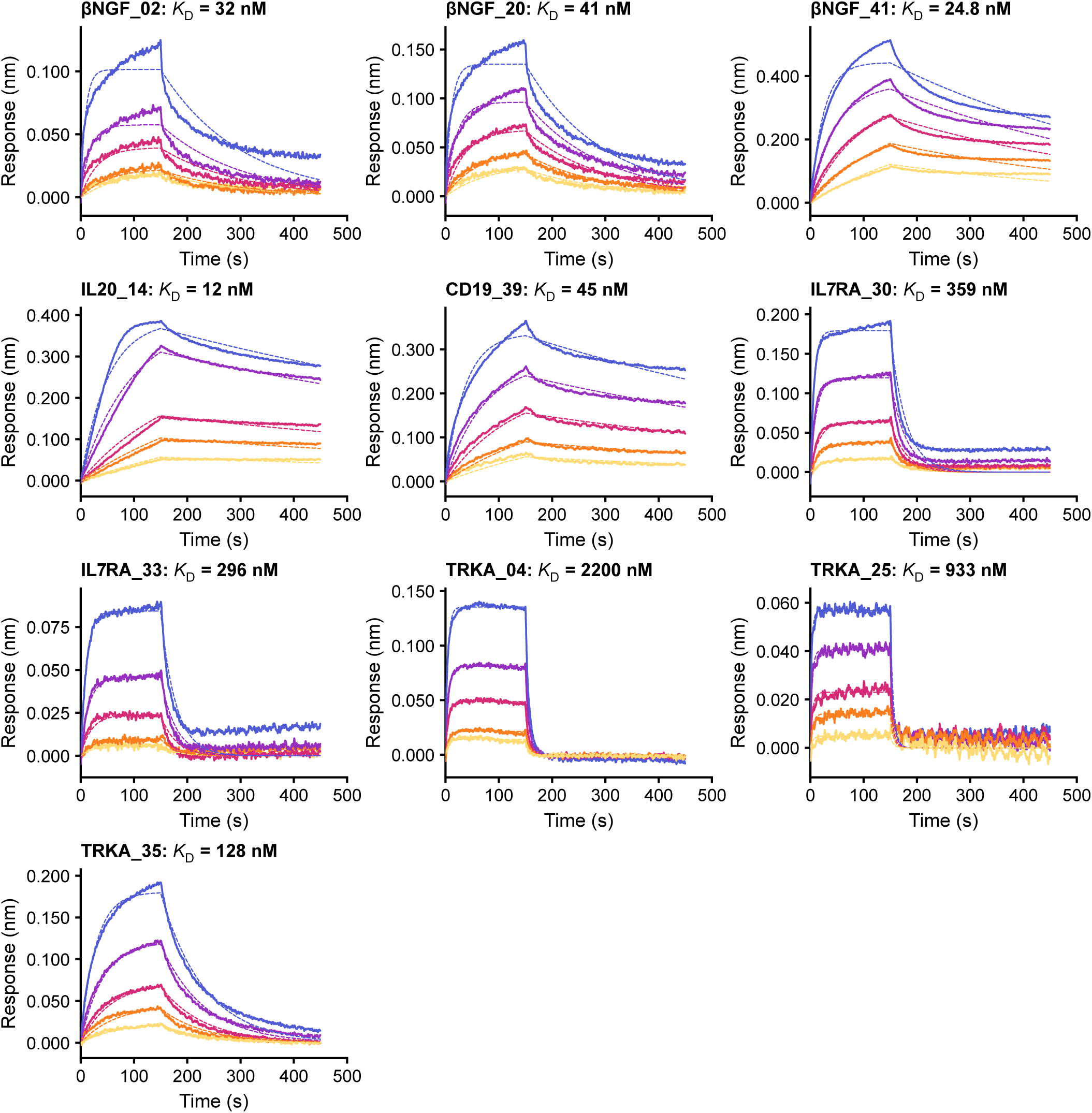
BLI binding analysis of antibodies against their respective recombinant antigens. Representative sensorgrams show concentration-dependent binding of antigen (β-NGF is a homodimer) at five 2-fold serial dilutions to immobilized antibody. βNGF_02, βNGF_20, and βNGF_41 were tested with antigen titrations starting at 500 nM; IL20_14 and TRKA_35 starting at 250 nM; and CD19_39, IL7RA_30, IL7RA_33, TRKA_04 and TRKA_25 starting at 1000 nM. Discrepancies between some experimental traces and their corresponding fits, for example the multiphasic profiles observed for βNGF_02 and βNGF_20, suggest non-ideal binding behavior, likely reflecting a component of nonspecific binding and/or avidity effects associated with contaminating aggregates.

**Figure S10.**
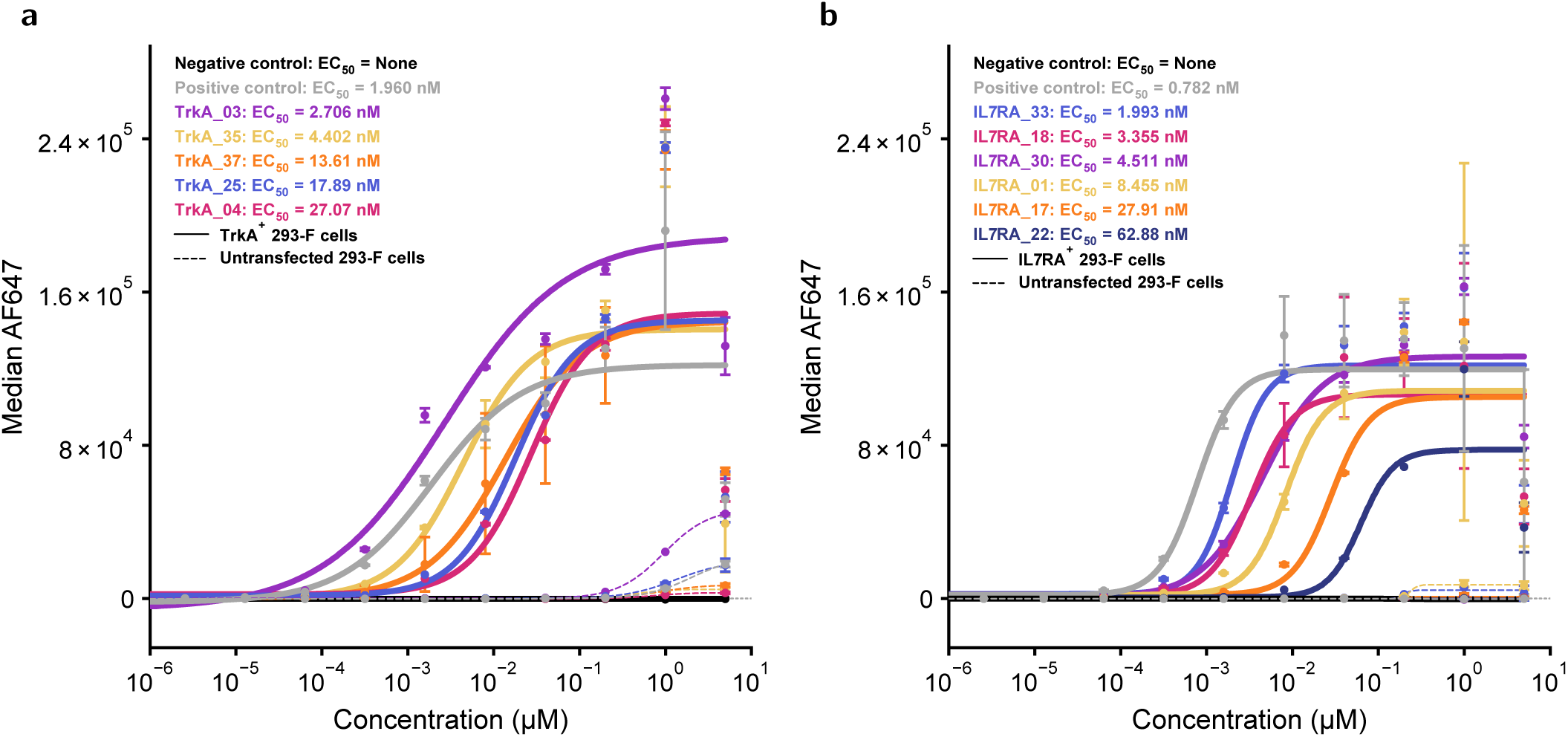
Cell binding profile of TRKA and IL7RA primary hits. (a) Concentration-dependent binding of selected antibodies to HEK293-F cells transfected with TRKA-containing plasmid (solid lines) or to untransfected cells (dashed lines) was measured by flow cytometry. Binding responses were fit using nonlinear regression to determine on-cell EC_50_ values. (b) Similar experiments were run for IL7RA primary hits with IL7RA-transfected HEK293-F cells (solid lines) or untransfected HEK293-F controls (dashed lines).

**Figure S11.**
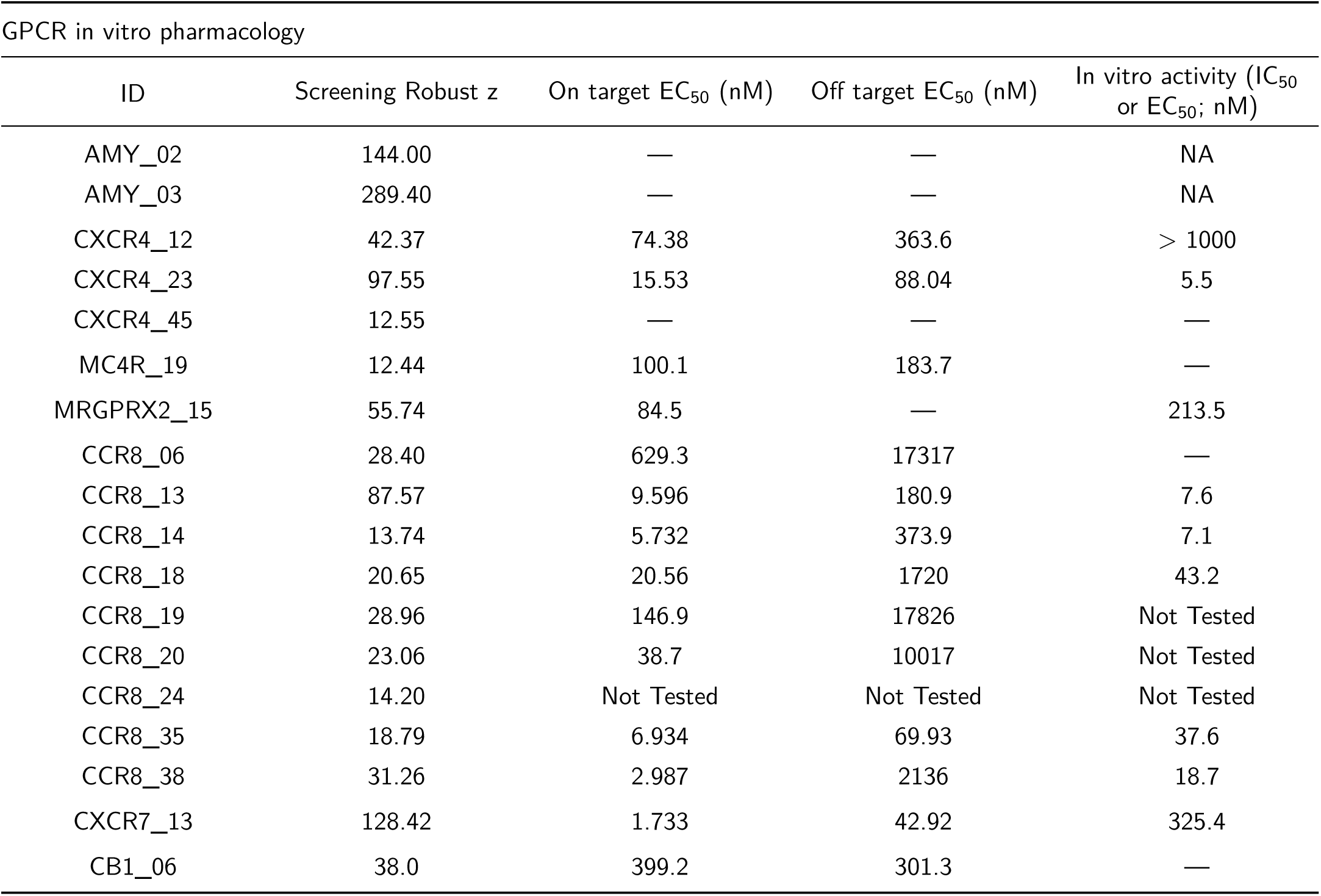
***In vitro*** pharmacological characterization of GPCR-targeting antibody hits. Summary of screening activity and functional characterization of selected GPCR antibody hits. Screening Robust z-scores are shown alongside EC_50_ values for on-target and off-target activity and potency (IC_50_ or EC_50_) in target-specific *in vitro* functional assays. An em dash, NA, and Not Tested indicate no detectable binding or activity, not applicable, and not tested, respectively. The value of 325.4 nM for CXCR7_13, which was tested in agonist and antagonist assay formats, refers to antagonist activity.

**Figure S12.**
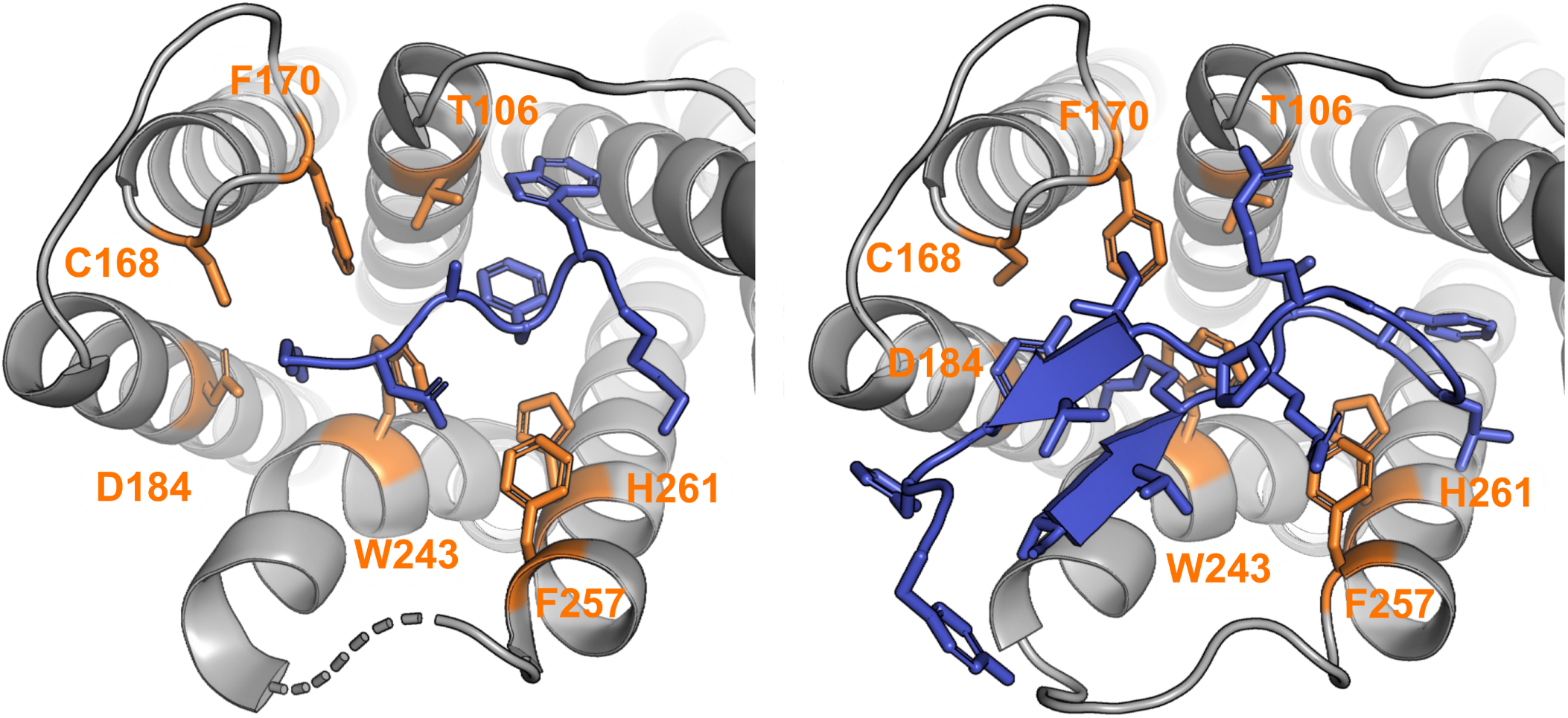
Structure of MRGPRX2 bound to cortistatin-14 (PDB 7S8L; left) compared with the MRGPRX2_15 HCDR3 binding pose (right). MRGPRX2 is shown in grey, cortistatin and the MRGPRX2_15 HCDR3 in blue, and receptor residues within 4 Å of both ligands are highlighted in orange.

## Supplementary tables

**Supplementary Table S1.**
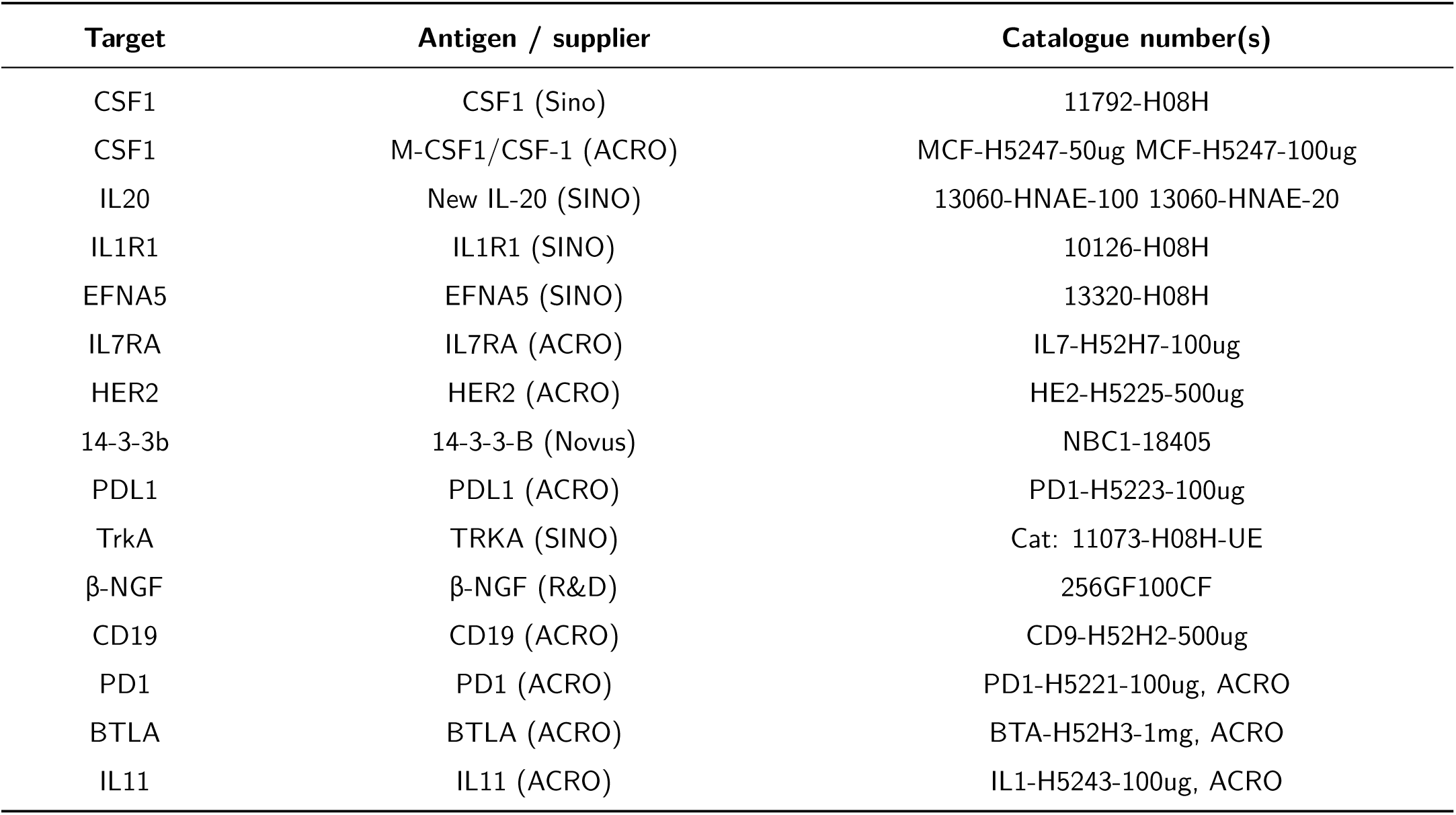
Recombinant antigens and commercial sources. Supplier descriptions and catalogue identifiers are reproduced from the supplied reagent list. The two CSF1 entries represent separate source products.

**Supplementary Table S2.**
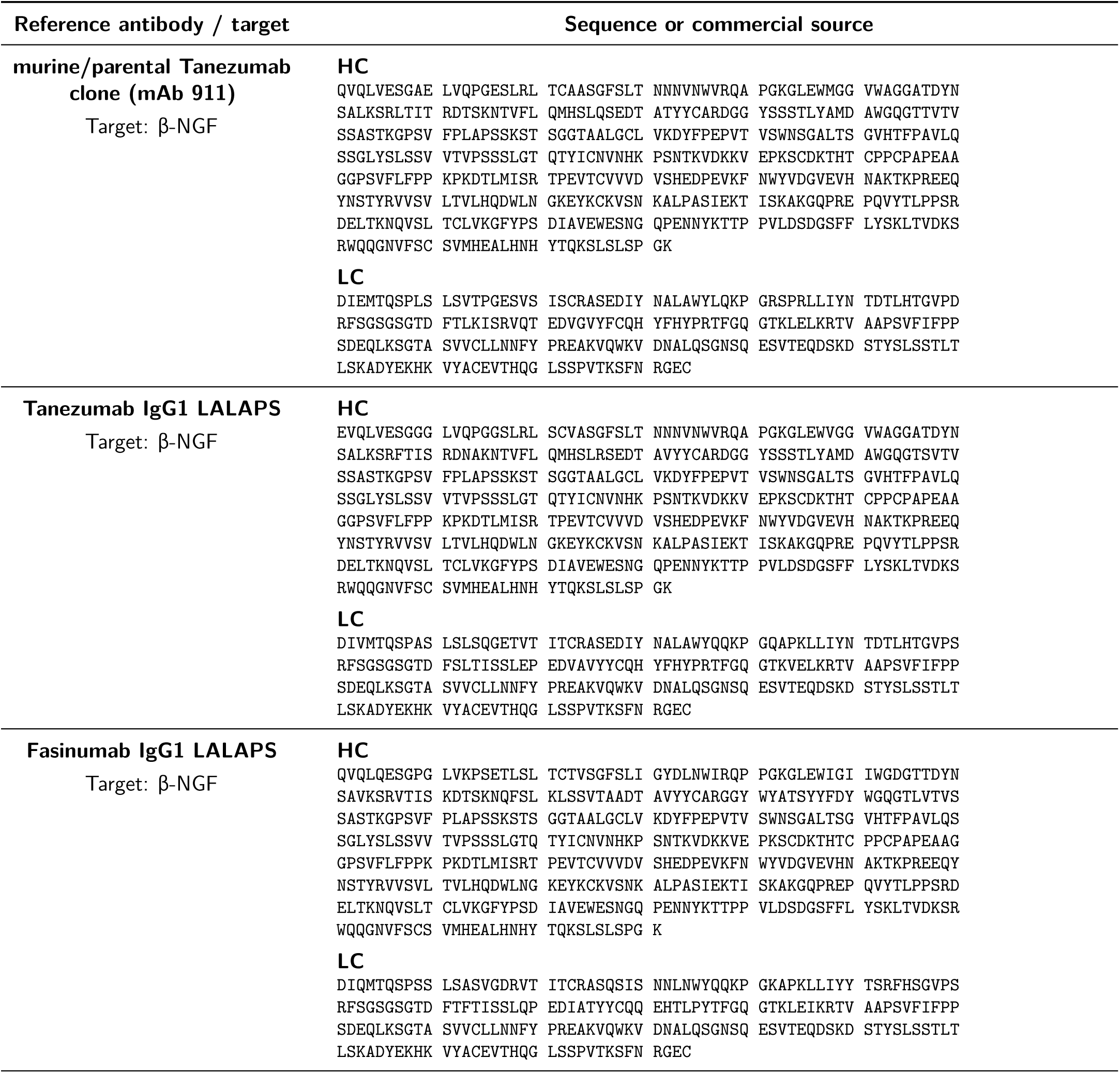
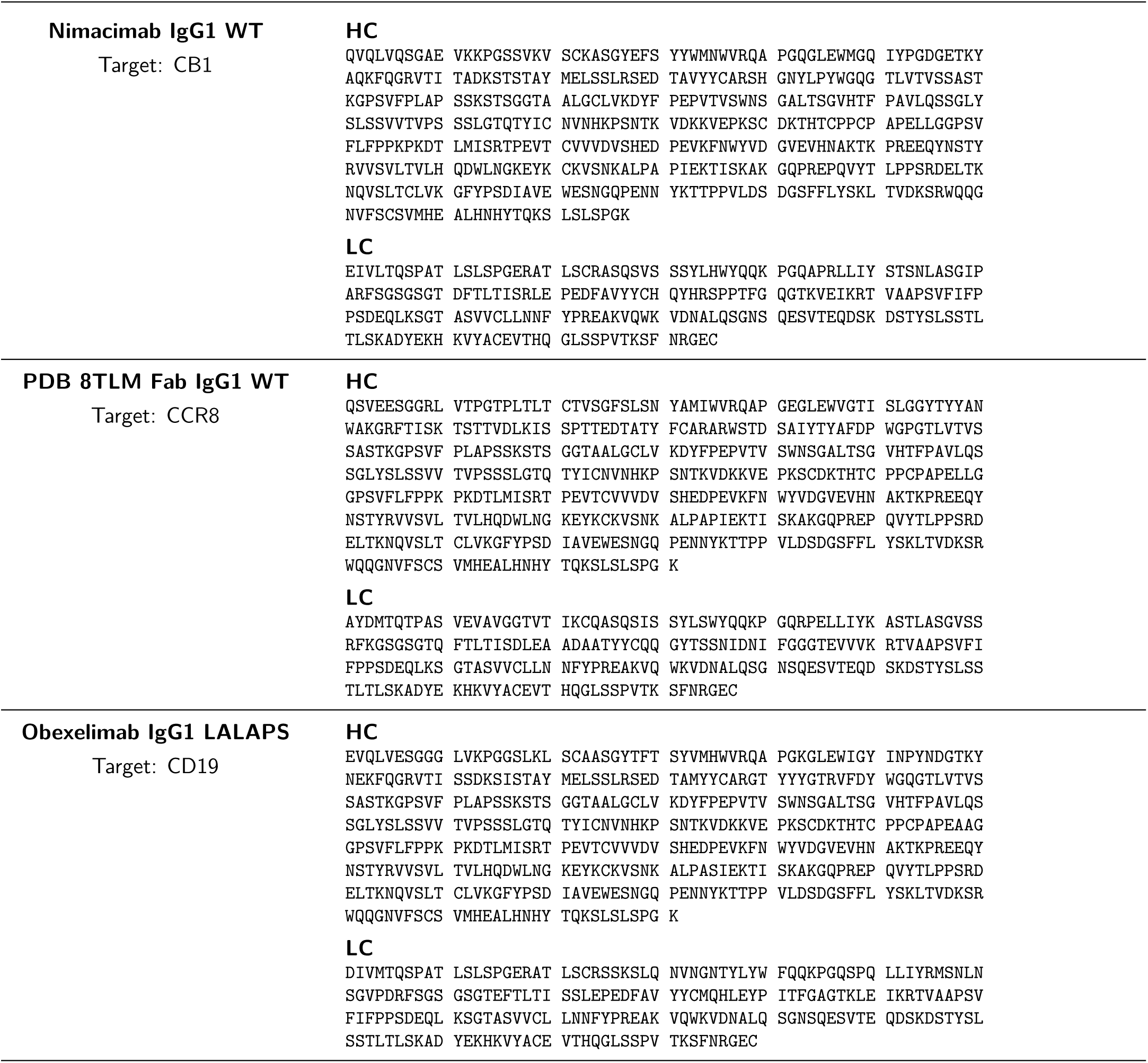
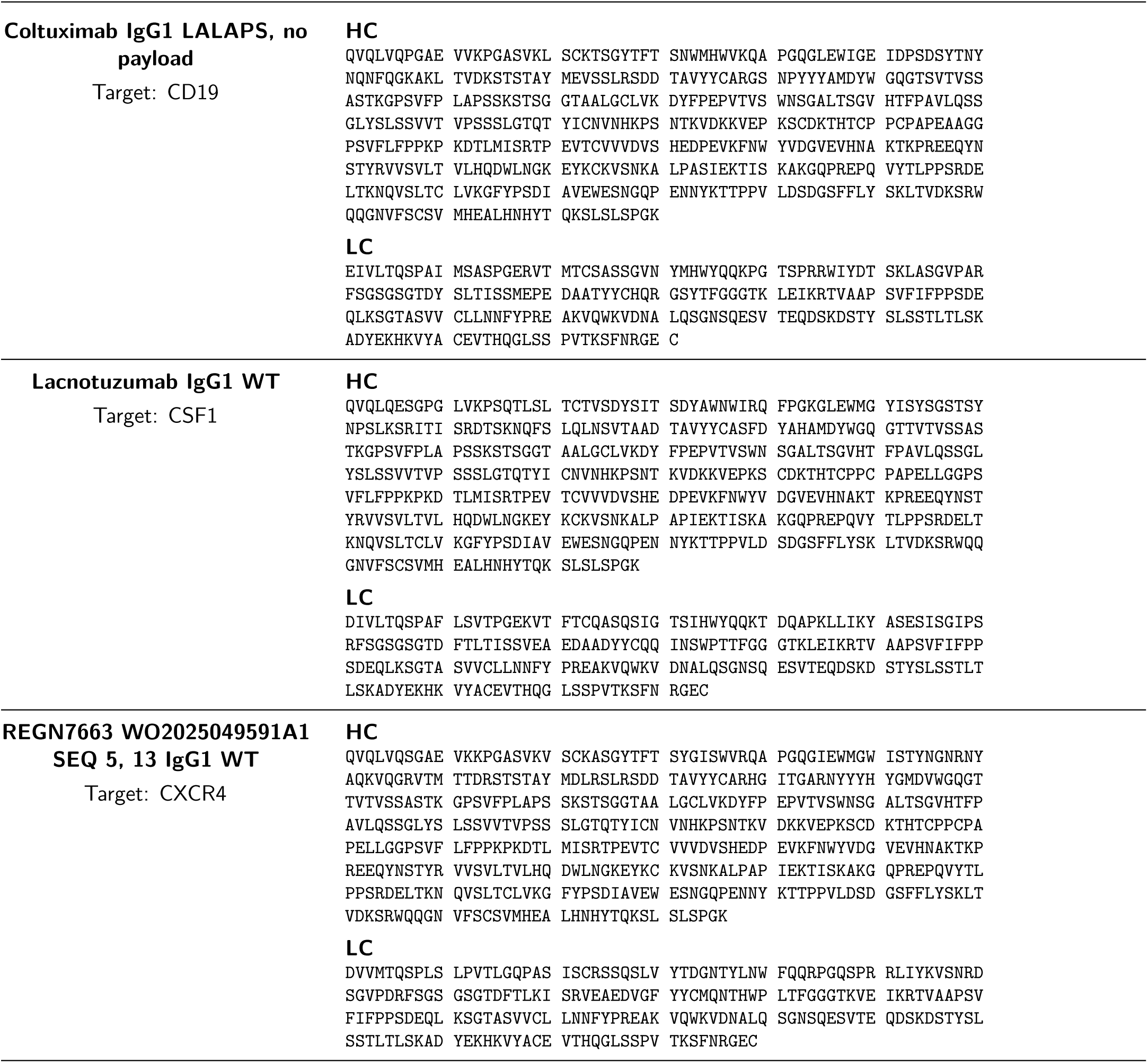
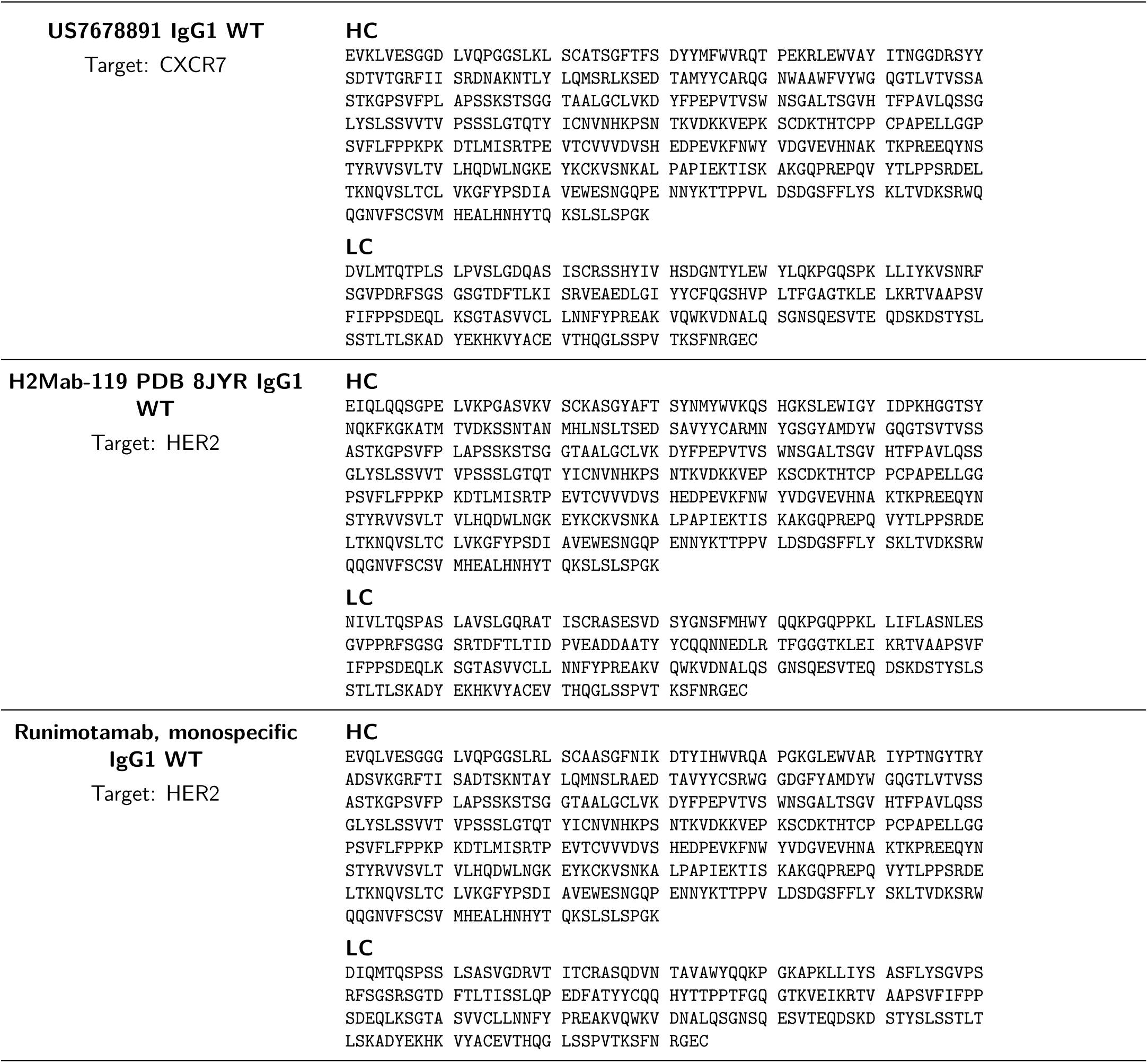
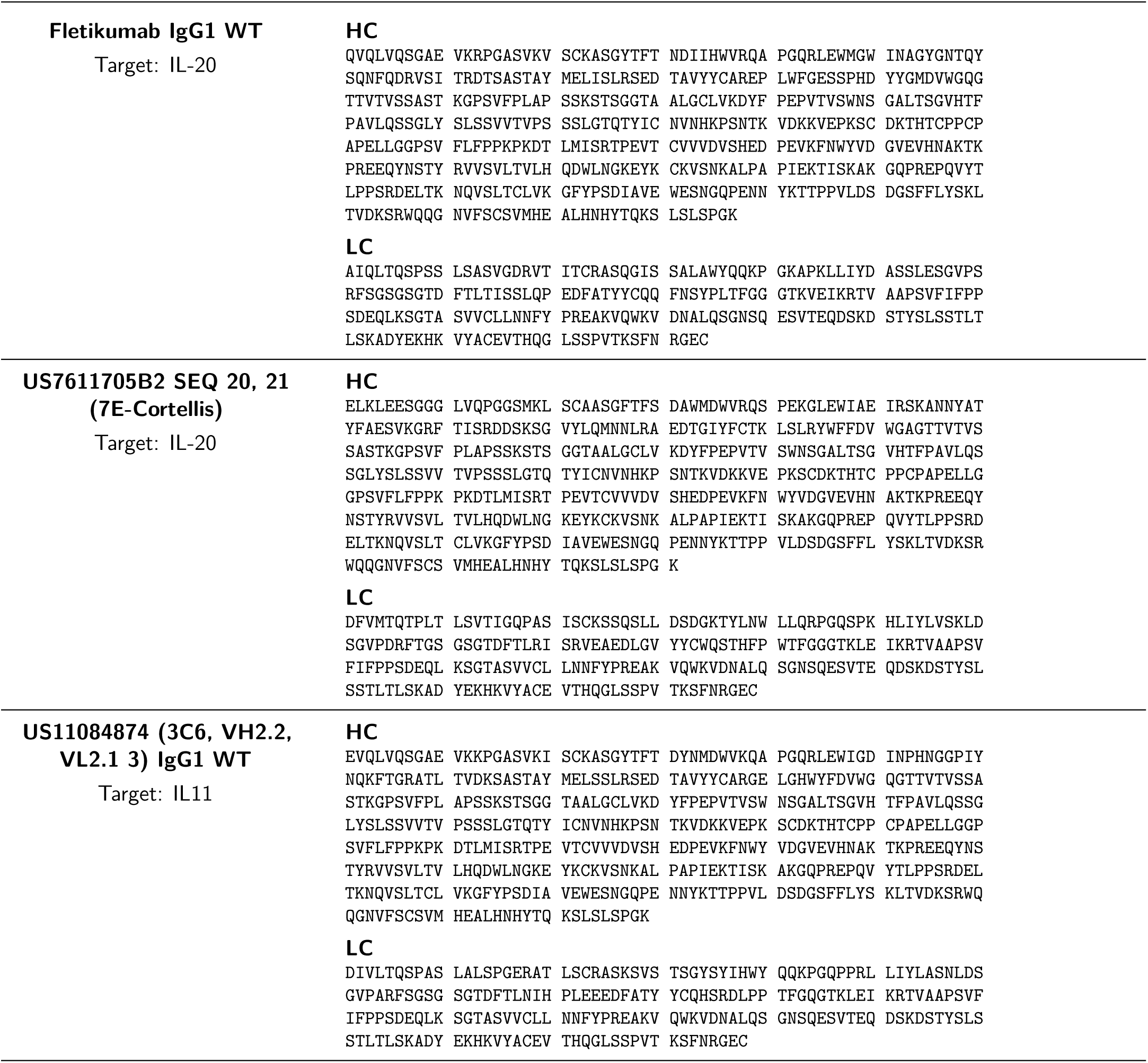
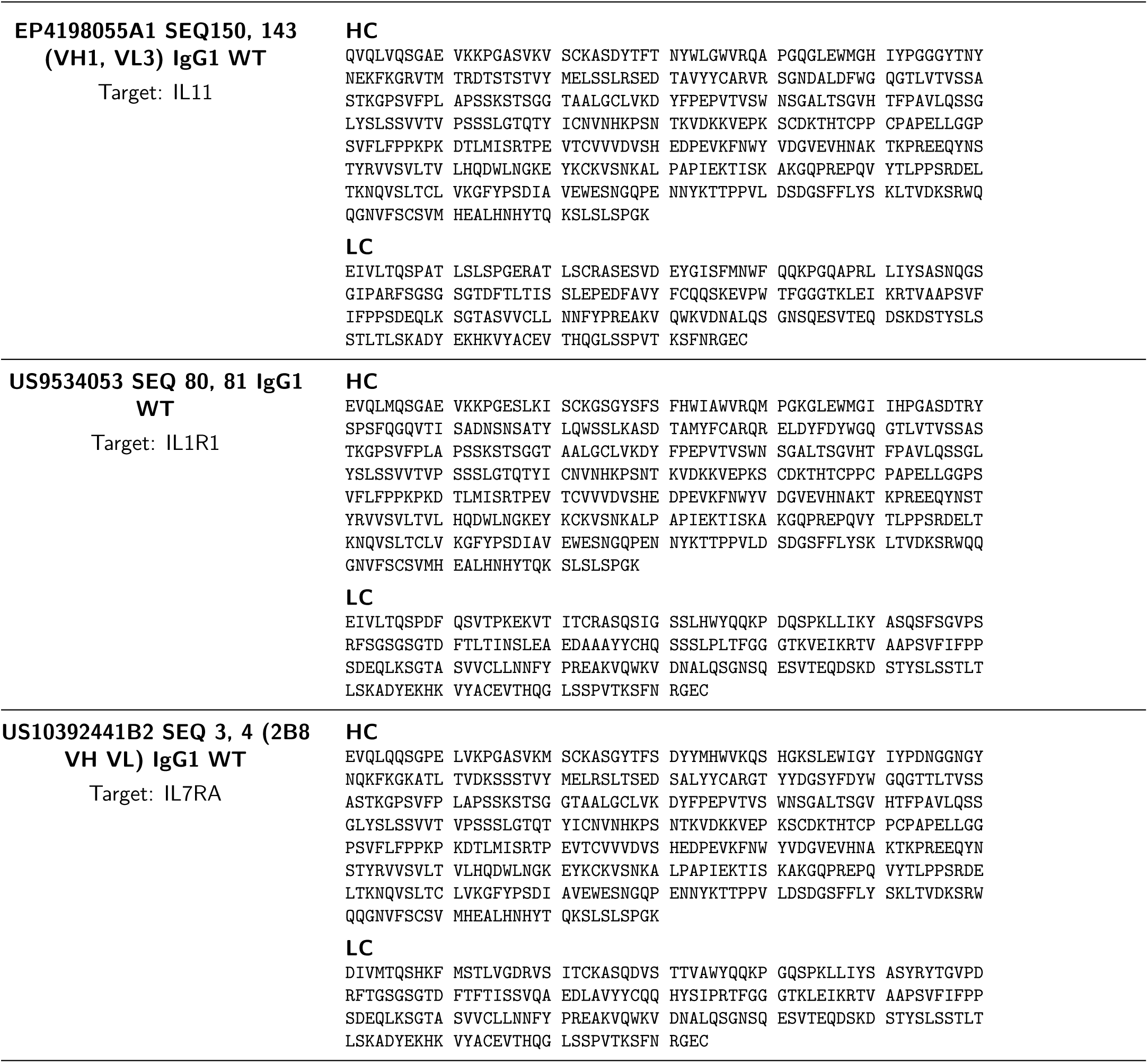
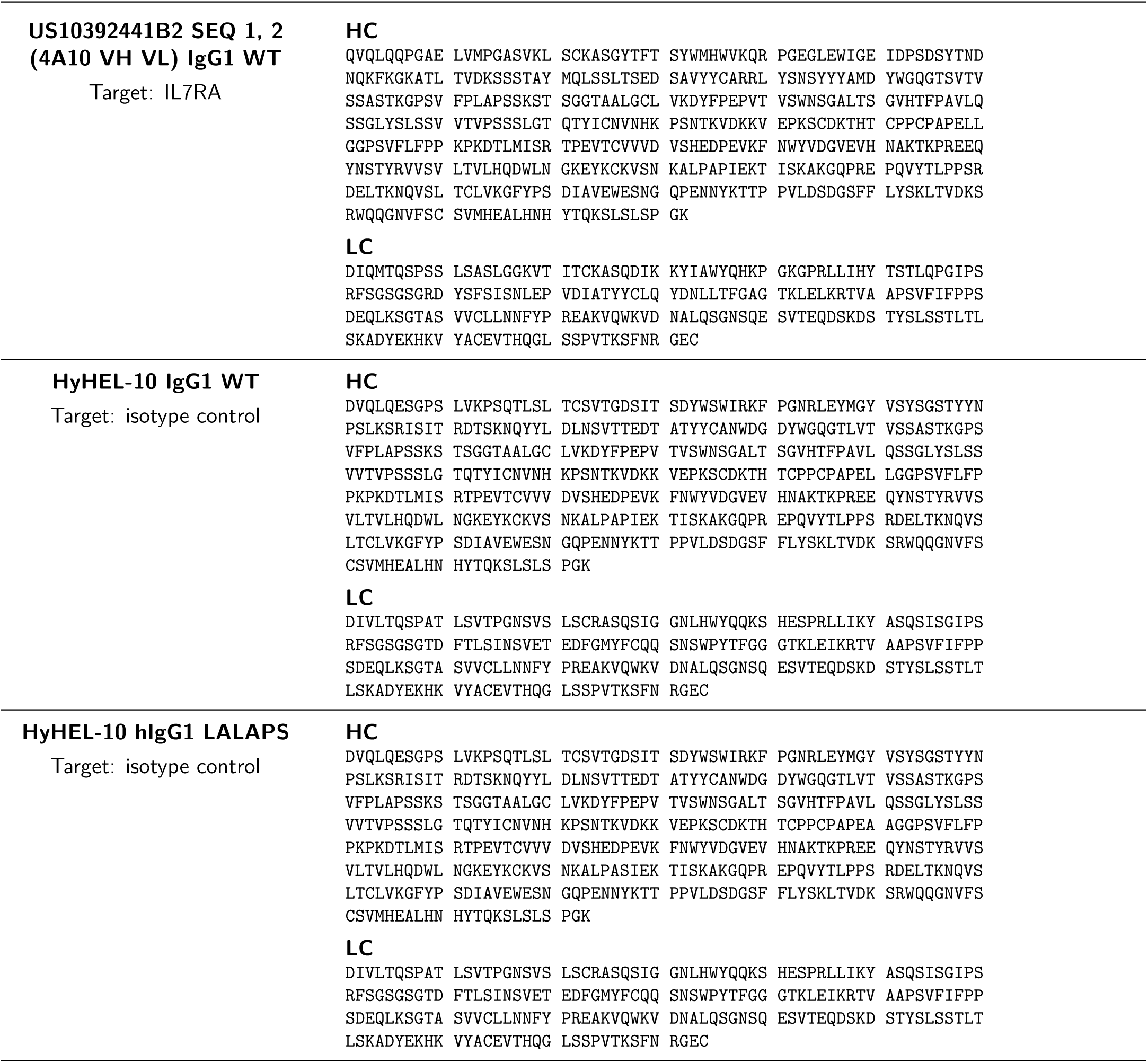
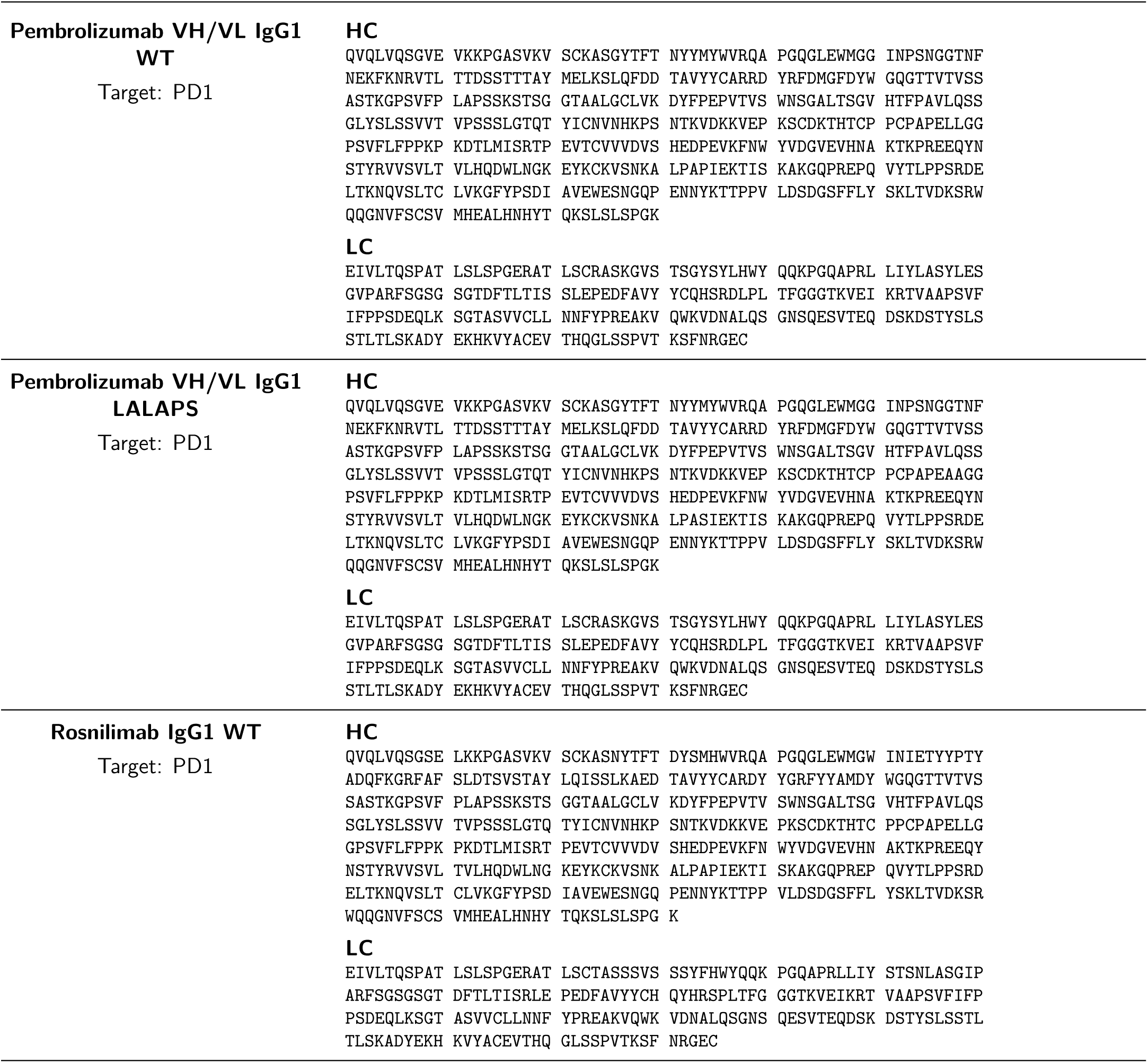
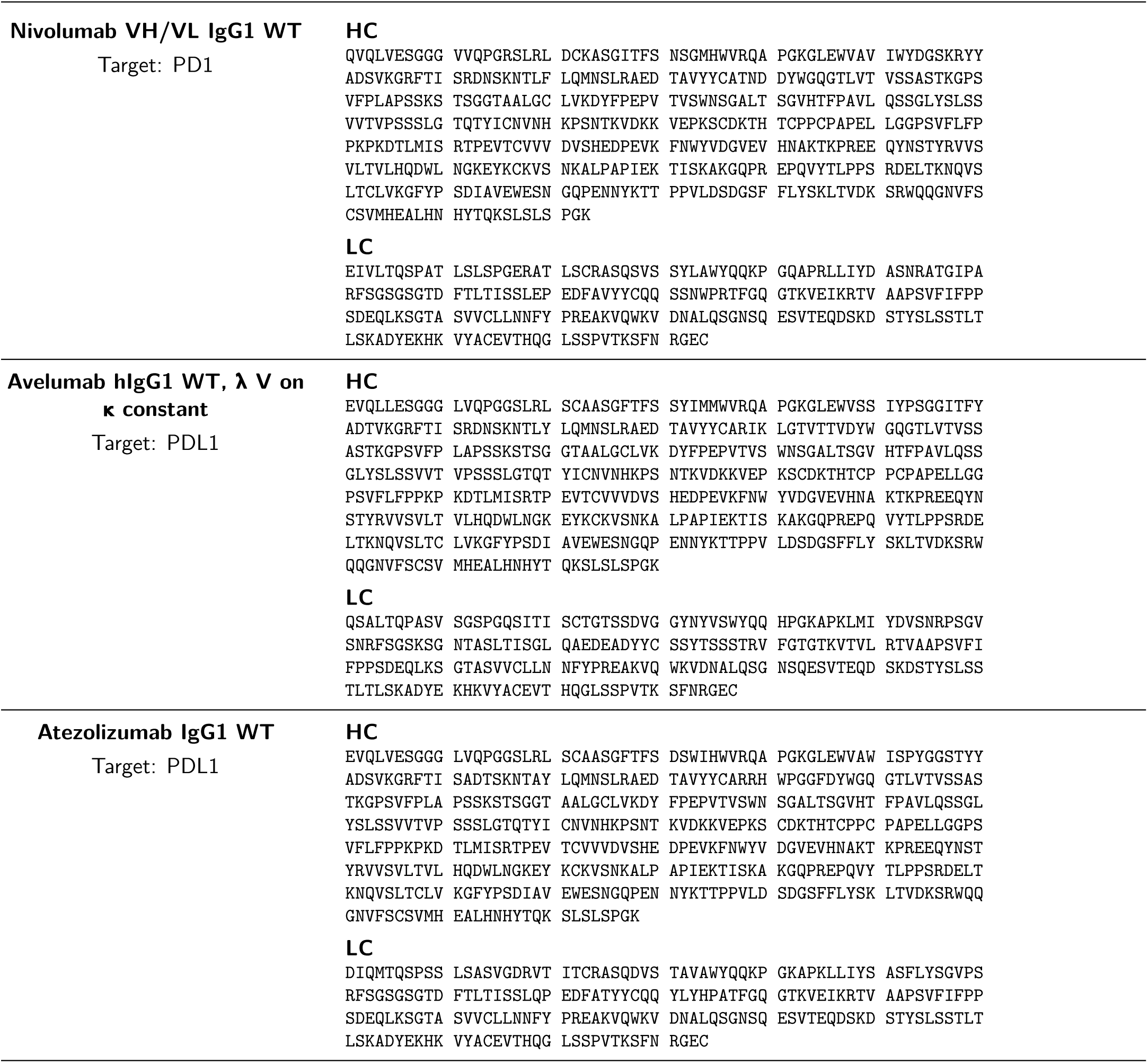
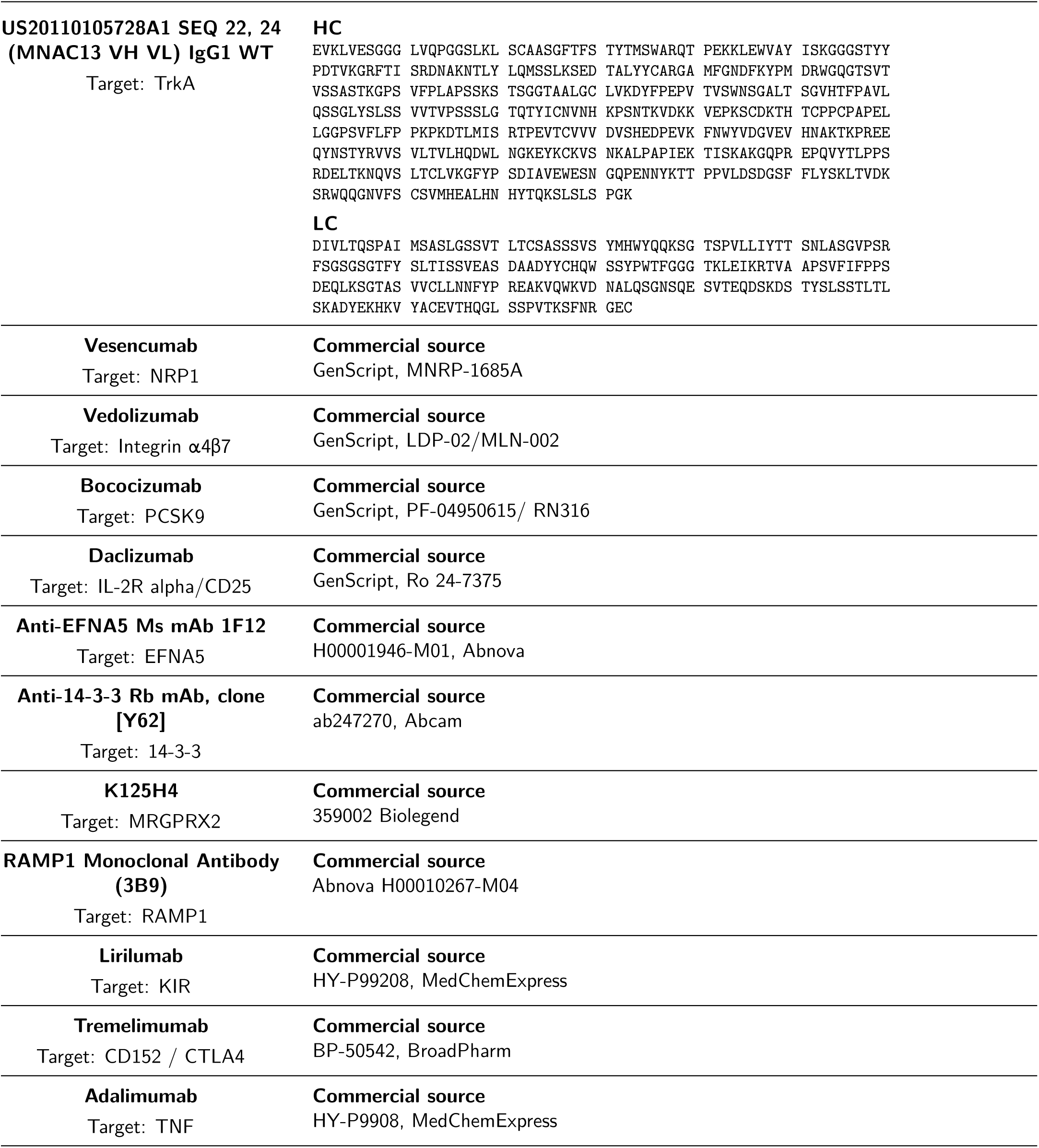
Reference antibodies: chain sequences and commercial sources. HC, heavy chain; LC, light chain. Sequences are reproduced without residue changes; spaces separate blocks of ten residues. For purchased antibodies, the supplied commercial source replaces the sequence fields.

